# Uncoupling the TFIIH Core and Kinase Modules Leads To Misregulated RNA Polymerase II CTD Serine 5 Phosphorylation

**DOI:** 10.1101/2023.09.11.557269

**Authors:** Gabriela Giordano, Robin Buratowski, Célia Jeronimo, Christian Poitras, François Robert, Stephen Buratowski

## Abstract

TFIIH is an essential transcription initiation factor for RNA polymerase II (RNApII). This multi-subunit complex comprises two modules that are physically linked in S. cerevisiae by the subunit Tfb3 (MAT1 in metazoans). The Core Module, with two DNA-dependent ATPases and several additional subunits, promotes DNA unwinding. The Kinase Module phosphorylates the C-terminal domain (CTD) of RNApII subunit Rpb1, initiating a cycle of CTD modifications that coordinate exchange of initiation and elongation factors. Why these two disparate activities are bundled into one factor is not obvious, but the connection may provide temporal coordination during early initiation. When Tfb3 is split into two parts to uncouple the TFIIH modules, the resulting cells are viable but grow very slowly. Chromatin immunoprecipitation of the split TFIIH shows that the Core Module, but not the Kinase, is properly recruited to promoters. Instead of the normal promoter-proximal peak, high CTD Serine 5 phosphorylation is seen throughout transcribed regions. Therefore, coupling the TFIIH modules is necessary to localize and limit CTD kinase activity to early stages of transcription. These results are consistent with the idea that the two TFIIH modules began as independent functional entities that later became connected by Tfb3 during early eukaryotic evolution.

**Impact Statement:** The TFIIH subunit Tfb3/MAT1 can be split into two parts to uncouple the TFIIH kinase and DNA translocase modules, resulting in unfocused CTD phosphorylation.

## INTRODUCTION

The three nuclear RNA polymerases in eukaryotes share many features, reflecting their evolution from a single precursor (reviewed in (Vannini & Cramer, 2012)). Their multiple subunits are either homologous or identical, and the initiation factors needed to direct all three polymerases to their appropriate target promoters include both the TATA-Binding Protein (TBP) and a TFIIB-like factor. Extending the parallels, RNA polymerases I and III contain intrinsic subunits that resemble the RNA polymerase II (RNApII) initiation factors TFIIE or TFIIF, which help direct the template DNA into the active site cleft.

In contrast to this overall mechanistic conservation, RNApII initiation factor TFIIH has no obvious counterparts in the other polymerase systems. As the final basal factor to assemble into the Pre-Initiation Complex (PIC), TFIIH performs two functions that are unique to RNApII (Thomas & Chiang, 2006). The first is the use of ATP hydrolysis to accelerate promoter melting, and the second is phosphorylation of a unique C-terminal domain (CTD) on Rpb1, the largest RNApII subunit. These two activities presumably evolved after the divergence of the three RNA polymerases.

The two TFIIH functions map to distinct sub-complexes, sometimes called modules, that are apparent in the TFIIH structure (**Figure 1A**) (He et al., 2016; Robinson et al., 2016; Schilbach et al., 2017). Ssl2 (XPB in metazoans) is the DNA-dependent ATPase that pushes double-stranded DNA towards the RNApII active site, leading to strand separation and placement of the template strand in the polymerase active site (Grünberg et al., 2012). Ssl2/XPB interacts with DNA downstream of RNApII, held in place by six other subunits (Tfb1, Tfb2, Tfb4, Tfb5, Rad3/XPD, and Ssl1) within the “Core” TFIIH subcomplex. TFIIH binding to the PIC is mediated by interactions of the Core with TFIIE and the Rpb4/7 stalk of RNApII (He et al., 2016; Robinson et al., 2016; Schilbach et al., 2017). Interestingly, Rad3 (XPD in metazoans) is a second DNA-dependent ATPase within the core, but with opposite DNA polarity relative to Ssl2/XPB. Surprisingly, Rad3/XPD makes no contact with DNA in PIC structures, and catalytically inactive Rad3 mutants do not affect transcription (Sung et al., 1988; Winkler et al., 2000). It is not obvious why there are so many Core TFIIH subunits with no clear contribution to the promoter melting function other than positioning Ssl2/XPB. One relevant consideration is that TFIIH also functions as a DNA nucleotide excision repair (NER) enzyme. In that process, both TFIIH ATPases unwind DNA to expose a short stretch of the damaged strand for lesion removal (Kim et al., 2023; van Eeuwen, Shim, et al., 2021; Yu et al., 2023). It seems plausible that the TFIIH Core module originally evolved for DNA repair, and only later was co-opted for RNApII transcription.

**Figure 1.**
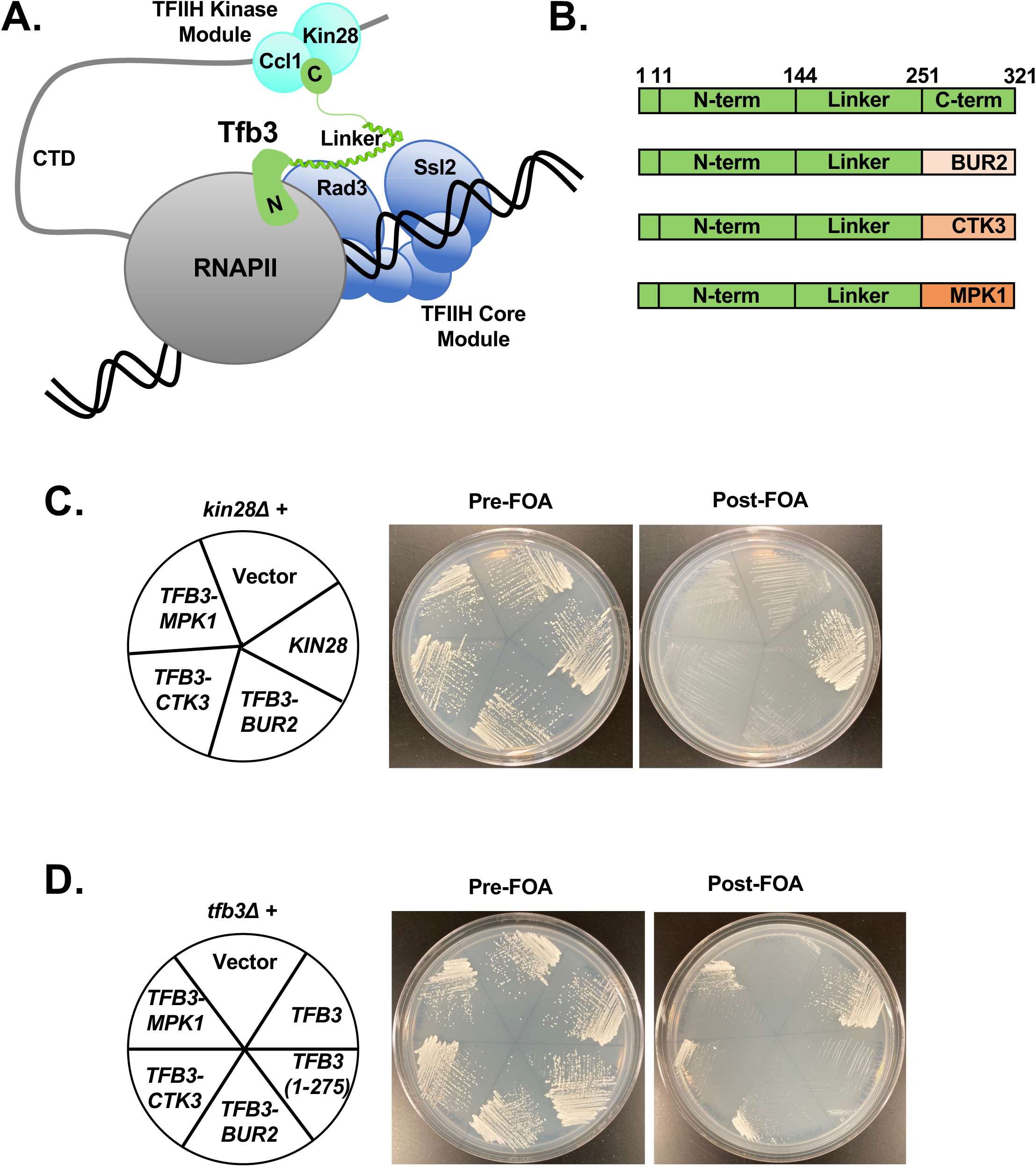
Tfb3 N-term-Linker fusions to other kinase modules can replace Tfb3, but not Kin28. **(A)** Schematic of TFIIH structure within the PIC. The domain structure of Tfb3 (green) is shown, connecting the TFIIH Kinase (cyan) and Core (blue) Modules. **(B)** Schematic of Tfb3 fusions to other kinase modules. **(C)** The *kin28Δ* strain YSB744 was transformed with pRS425 (Vector), pRS415-*KIN28*, pRS425-*TFB3-BUR2*, pRS425-*TFB3-CTK3*, or pRS425-*TFB3-MPK1*. Transformants were tested for the ability to replace a *KIN28/URA3* construct by plasmid shuffling. Cells were streaked on -LEU-TRP (center) or -LEU-TRP+5-FOA (right) plates for 3 days at 30℃. **(D)** pRS315 (Vector), pRS425-*TFB3*, pRS315/TFB3Δ2 (aa 1-275) (Feaver et al., 2000), pRS425-*TFB3-BUR2*, pRS425-*TFB3-CTK3*, and pRS425-*TFB3-MPK1* were transformed into *tfb3Δ* strain SHY907 (Warfield et al., 2016), replacing the plasmid-expressed wild-type gene by plasmid shuffling. Streaks shown were grown on –LEU for 4 days (center) or on -LEU+5-FOA plates (right) for 9 days.

The second TFIIH sub-complex carries the CTD phosphorylation activity. The Rpb1 CTD is made up of multiple repeats of the sequence YSPTSPS, and its changing phosphorylation patterns help recruit various mRNA processing factors during different stages of the transcription cycle (Buratowski, 2009). The Kin28 (CDK7 in metazoans) kinase and its cyclin partner Ccl1 (Cyclin H) together phosphorylate CTD Serine 5 at the promoter. PIC structures show that the Mediator complex helps guide the CTD to the kinase module (Robinson et al., 2016; Schier & Taatjes, 2021; Schilbach et al., 2017; Schilbach et al., 2023). CTD phosphorylation triggers release of Mediator, as well as subsequent recruitment of mRNA capping enzyme and other factors that function near 5’ ends of mRNAs (reviewed in (Buratowski, 2009)). Like the Core, the Kinase module also has a second function unrelated to transcription. At least in metazoans, CDK7/cyclin H serves as a CDK-activating kinase (CAK), phosphorylating the T-loop of other cyclin-dependent kinases to promote an active conformation (Fisher & Morgan, 1994). This is not a universal function, as *Saccharomyces cerevisiae* and some other species have a separate CAK enzyme (Cismowski et al., 1995; Espinoza et al., 1996; Kaldis et al., 1996).

Nonetheless, it seems likely that the CDK7 activity targeting RNApII evolved from a pre-existing CDK that had either CAK activity or some other function.

The two TFIIH subcomplexes are linked by the Tfb3 subunit (known as MAT1 in metazoans). Tfb3/MAT1 consists of two globular domains and a linker region. The N-terminal domain contains a zinc-binding RING finger and is part of the Core module, where it helps incorporate TFIIH into the PIC through direct interactions with RNApII and TFIIE (Schier & Taatjes, 2021). The Tfb3 C-terminal domain contacts both the kinase and cyclin subunits, and this interaction promotes an active kinase conformation (Greber et al., 2020; Peissert et al., 2020; van Eeuwen, Li, et al., 2021). In many TFIIH and PIC structures the linker is not visible, presumably due to flexibility. However, when it is seen (Abril-Garrido et al., 2023; Greber et al., 2019), the linker emerges from the N-terminal domain as a long alpha-helix running along the interface between the two ATPase subunits, followed by a turn and a short stretch of helix just N-terminal to a disordered region that connects to the C-terminal region (see schematic in **Figure 1A**). A deletion study in yeast showed that Tfb3 tolerates 30 amino acid deletions throughout the linker, but that the N- and C-terminal domains were essential for viability (Warfield et al., 2016).

Given its modular structure, we tested whether other CTD kinases could be fused to Tfb3 to bypass the requirement for Kin28/CDK7. While the results have so far been negative, we unexpectedly found that cells lacking the C-terminal domain of Tfb3 were actually viable, albeit very slow growing. Remarkably, growth is significantly rescued when the two parts of Tfb3 are expressed as separate proteins. In these cells, the TFIIH kinase module is no longer tethered to the Core TFIIH. ChIP experiments show that the Core Module, but not the Kinase, is still promoter localized. Surprisingly, phosphorylation of CTD Serine 5 is now seen at higher than normal levels throughout transcribed regions, indicating that the untethered kinase accesses RNApII throughout the cell cycle. These results are consistent with the idea that the two TFIIH modules began as independent entities that only later became fused into a single complex to provide coordinated regulation of the TFIIH kinase and promoter melting activities.

## RESULTS

### Tethering other kinase modules to Tfb3 does not bypass the requirement for Kin28

Given the apparent modularity of TFIIH, we asked whether tethering other kinases to Tfb3 could functionally replace Kin28. Although Kin28 phosphorylates CTD Serine 5 in vivo, other kinases can phosphorylate this residue in vitro. Two other transcription kinase complexes, Bur1/Bur2 (the yeast homologs of CDK9/cyclin T) and Ctk1/Ctk2/Ctk3 (yeast CDK12/Cyclin K), phosphorylate the Rpb1 linker (Chun et al., 2019) and CTD Serine 2 (Cho et al., 2001), respectively. However, both are reported to also have activity on CTD Serine 5, at least in vitro (Corden, 2013; Eick & Geyer, 2013; Glover-Cutter et al., 2009). Accordingly, we constructed Tfb3 fusions **(Figure 1B, Supplemental Table 1)** in which the C-terminal region (C-term) was replaced with either Bur2 (the cyclin partner of Bur1) or Ctk3 (the third subunit within the Ctk1/Ctk2 CDK/cyclin complex). We also created a fusion of the Tfb3 N-terminal domain (N-term) and linker to Mpk1, a single subunit kinase also reported to have activity on CTD Serine 5, as well as Ser2 and Tyr1 (Yurko et al., 2017).

The three fusion constructs were introduced into a yeast strain lacking the chromosomal copy of *KIN28*. Expression of the fusion proteins was confirmed by immunoblotting (**Figure 1 – figure supplement 1**). None of the fusions supported viability after plasmid shuffling to remove *KIN28* (**Figure 1C**), indicating that tethering the other kinases could not substitute for Kin28 function. As a control experiment, the Tfb3 fusions were also tested for complementation of a *TFB3* deletion. Surprisingly, these shuffled strains proved to be viable, albeit so slowly growing that 5-7 days were needed to see colonies **(Figure 1D)**. Complementation of *tfb3Δ* indicates that the fusion proteins were incorporated into TFIIH. This result was unexpected because a *TFB3* allele encoding amino acids 1-275, and therefore lacking the C-term, did not support growth, consistent with earlier reports that the C-terminal domain is essential for Tfb3 function and cell viability (Feaver et al., 2000; Warfield et al., 2016).

### The N-terminal domain is the only essential part of Tfb3

The unexpected complementation of *tfb3Δ* by the Tfb3 fusions raised the question of whether the Tfb3 C-term was truly essential for viability. Earlier studies may not have waited long enough to see growth. Also, the restriction fragment deletion used to create the non-complementing Tfb3(1-275) allele used in **Figure 1D** also removed the 3’ untranslated region and polyA site (Feaver et al., 2000), which could further impair function beyond the loss of the C-term. Accordingly, we created a new set of Tfb3 truncations, each lacking either one or two of the three major Tfb3 domains **(Figure 2A)**, and used plasmid shuffling to introduce them into a *tfb3Δ* strain. Cells lacking the N-term failed to grow, no matter how long the plates were incubated **(Figure 2B**).

**Figure 2.**
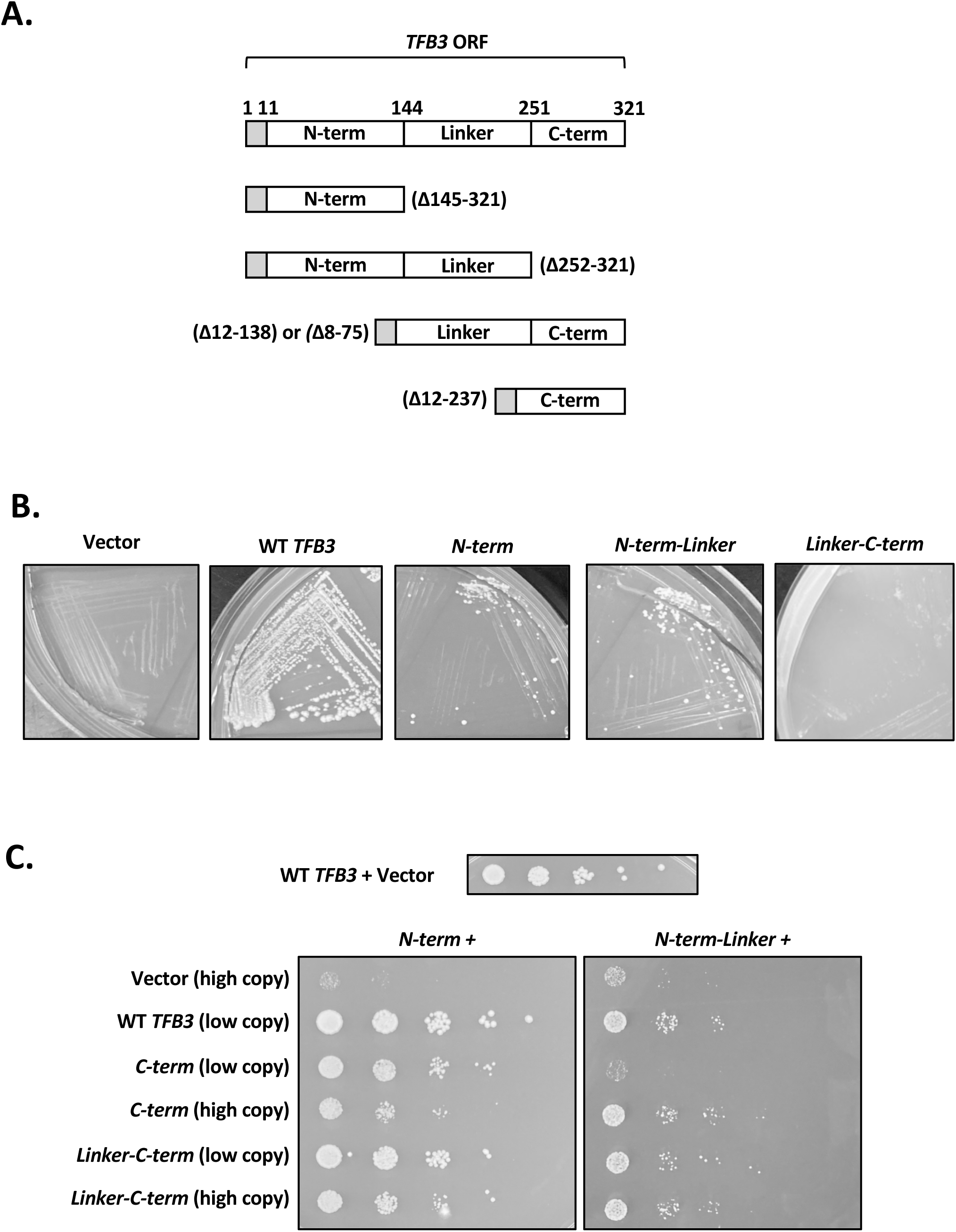
Tfb3 can be split into two functional parts. **(A)** Schematic of Tfb3 deletion constructs. Note that all constructs retained amino acids 1-11 at the N-terminus for ease of cloning and to avoid differential sensitivity to N-end rule degradation. **(B)** Plasmid shuffling was used to test pRS424 (Vector), pRS424-*TFB3* (WT *TFB3*), pRS424-*TFB3(*Δ145-321) (N-term), and pRS424-*TFB3* (Δ252-321) (N-term-Linker) for the ability to support growth of *tfb3Δ* shuffling strain SHY907/YF2456 (Warfield et al., 2016). Streaks shown were grown on -TRP+5-FOA plates for 5 days at 30℃ **(C)** Spot assay for growth. Tfb3 N-term (YSB3704) and N-term-Linker (YSB3722) strains were transformed with pRS425 (Vector), pSH1542 (WT *TFB3*), pRS315-TFB3(1-11, 238-Stop)-Flag1-TAP (C-term low copy), pRS425-TFB3(1-11, 238-Stop) (C-term high copy), pRS425-TFB3(1-11, 139-Stop) (Linker-C-term low copy), or pLH366 (Linker-C-term high copy). Ten-fold serial dilutions were spotted onto –LEU-TRP plates for 3 days. For comparison to normal growth, an isogenic strain carrying WT *TFB3* (YSB3788) was spotted in parallel (top strip).

In contrast, strains expressing only the N-term, or the N-term and Linker (N-term-Linker) domains, exhibited slow growth. These deletions grew at rates similar to the fusion proteins tested above, arguing that the fused kinase module did not provide an additional function or stability relative to a Tfb3 completely lacking any sequence C-terminal to the Linker region. We therefore conclude that, while important for normal growth, the C-term of Tfb3 (which is part of the Kinase Module) is not essential for viability.

### Expression of the TFIIH Core and Kinase Modules as independent complexes

Recent structural studies show that the C-terminal domain of Tfb3, or its metazoan homolog MAT1, stabilizes an active kinase conformation when bound to Kin28/CDK7 and its partner cyclin (Greber et al., 2020; Peissert et al., 2020; van Eeuwen, Li, et al., 2021). Therefore, the slow growth of strains lacking the Tfb3 C-terminal domain could be due to loss of kinase tethering to the Core Module, to lack of Kin28/CDK7 activation, or both.

To test the contribution of the Tfb3 kinase activation function separately from kinase tethering, parts of Tfb3 were expressed as two separate proteins (**Figure 2C**). Strains expressing either the N-term (left panel) or N-term-Linker (right panel) derivatives were transformed with plasmids expressing the indicated C-term constructs. These strains were tested for growth by serial dilution spotting, where the size of single colonies serves as a measure of relative growth rate. Again, strains lacking any C-term domain grew very slowly (Vector, top row). In the N-term strain (left panel), adding wild-type Tfb3 restored growth to normal. Co-expression of the separate N-term and C-term domains noticeably improved growth compared to N-term alone, despite the fact that this strain completely lacks the Linker region. Co-expression of the Linker-C-term construct rescued growth to an even greater extent, although not quite to wild-type rates. Therefore, the Linker does contribute to full function of the Kinase Module.

A similar pattern was seen for strains expressing the N-term-Linker construct (**Figure 2C**, right panel). Growth improvement was seen by co-expressing C-term, at least when a high copy plasmid is used to produce more protein. Co-expressing the Linker-C-term construct gave a similar growth improvement at both high and low copy, although no greater than the high copy C-term only construct. Altogether, the co-expression results show that the Tfb3 N-terminal and C-terminal domains can function independently of covalent physical linkage.

We considered the possibility that, despite the split Tfb3, the TFIIH Kinase and Core Modules could remain associated through other subunit contacts. To test this possibility, a tandem affinity purification (TAP) tag was used to precipitate full-length, C-term, or Linker-C-term Tfb3 derivatives using IgG beads. There was some variation in protein expression levels (**Figure 3A**, left panel, lanes 1-4), and reduced levels of the split Tfb3 may contribute to the slow growth phenotypes. However, similar amounts of all three TAP fusions were precipitated with IgG beads (lanes 5-8). The Kinase Module was intact, as subunits Kin28 and Ccl1 interacted with full-length or Tfb3 C-terminal constructs equally well (**Figure 3B**). An in vitro phosphorylation assay using GST-CTD as substrate showed that Kinase Modules from the truncated Tfb3 alleles retained activities similar to or slightly below that of full-length Tfb3 (**Figure 3 - figure supplement 1**).

**Figure 3.**
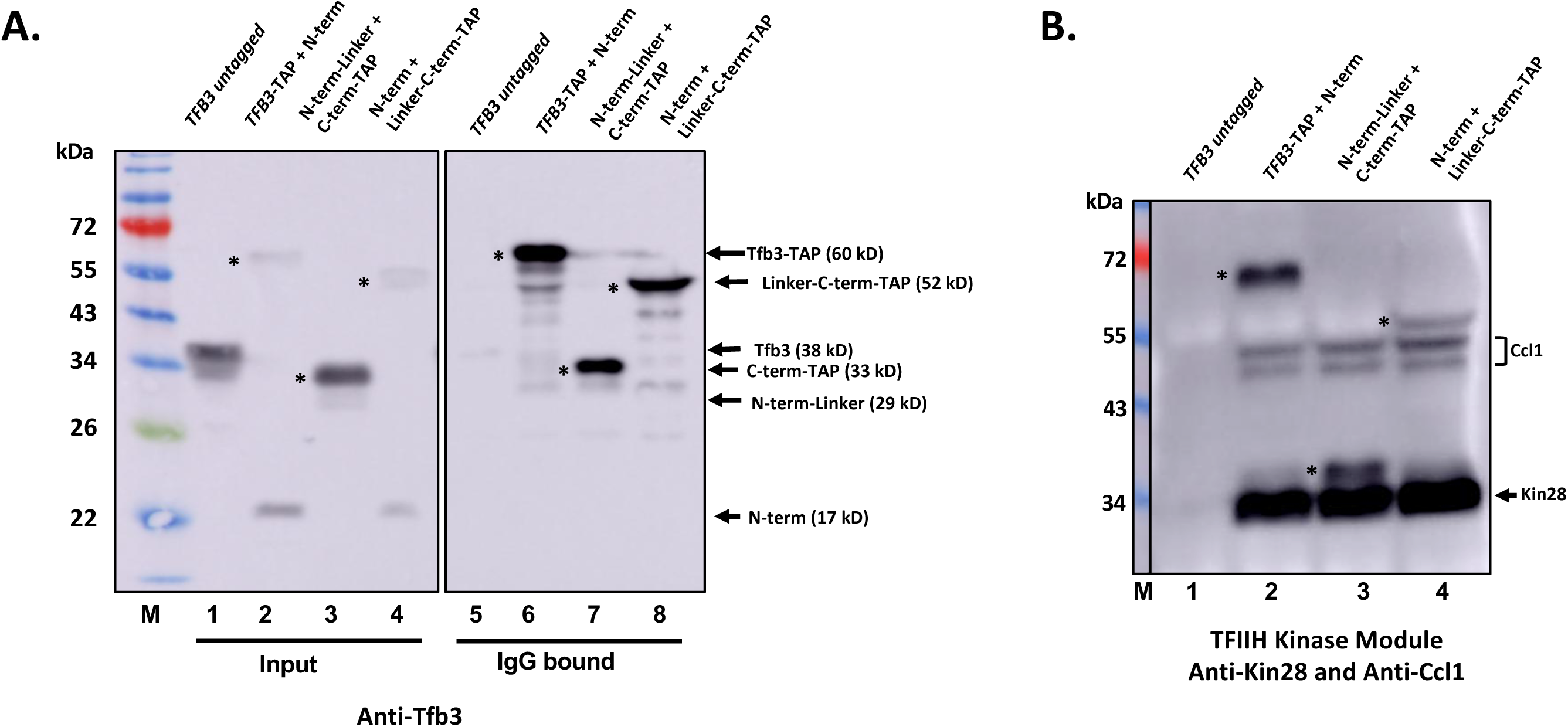
The TFIIH Kinase Module associates with the untethered Tfb3 C-terminal domain. **(A)** Whole cell extracts were made from YSB3788 (*TFB3* untagged), YSB3723 (*TFB3*-TAP + N-term), YSB3732 (N-term-Linker + C-term-TAP), and YSB3728 (N-term + Linker-C-term-TAP). Both Input extracts (lanes 1-4) and IgG-Agarose precipitated fractions (lanes 5-8) were resolved by SDS-PAGE and blotting for Tfb3. **(B)** To probe for other Kinase Module subunits, IgG precipitates were probed with anti-Ccl1 and anti-Kin28 antibodies. In all panels, asterisks mark TAP-tagged Tfb3 derivatives (which also react with secondary antibody due to the protein A component).

In contrast to the Kinase Module, the TFIIH Core Module (monitored by Core subunit Tfb1) no longer associates strongly with the uncoupled Tfb3 C-term derivatives (**Figure 4A**, compare lanes 3 and 4 to full-length Tfb3 in lane 2). The Linker-C-term derivative may retain some weak interaction with the Core, but the Tfb1 levels were just slightly above background binding in the untagged strain (lane 1), and far below that seen with full length Tfb3. We conclude that the split Tfb3 does indeed dissociate the two TFIIH modules.

**Figure 4.**
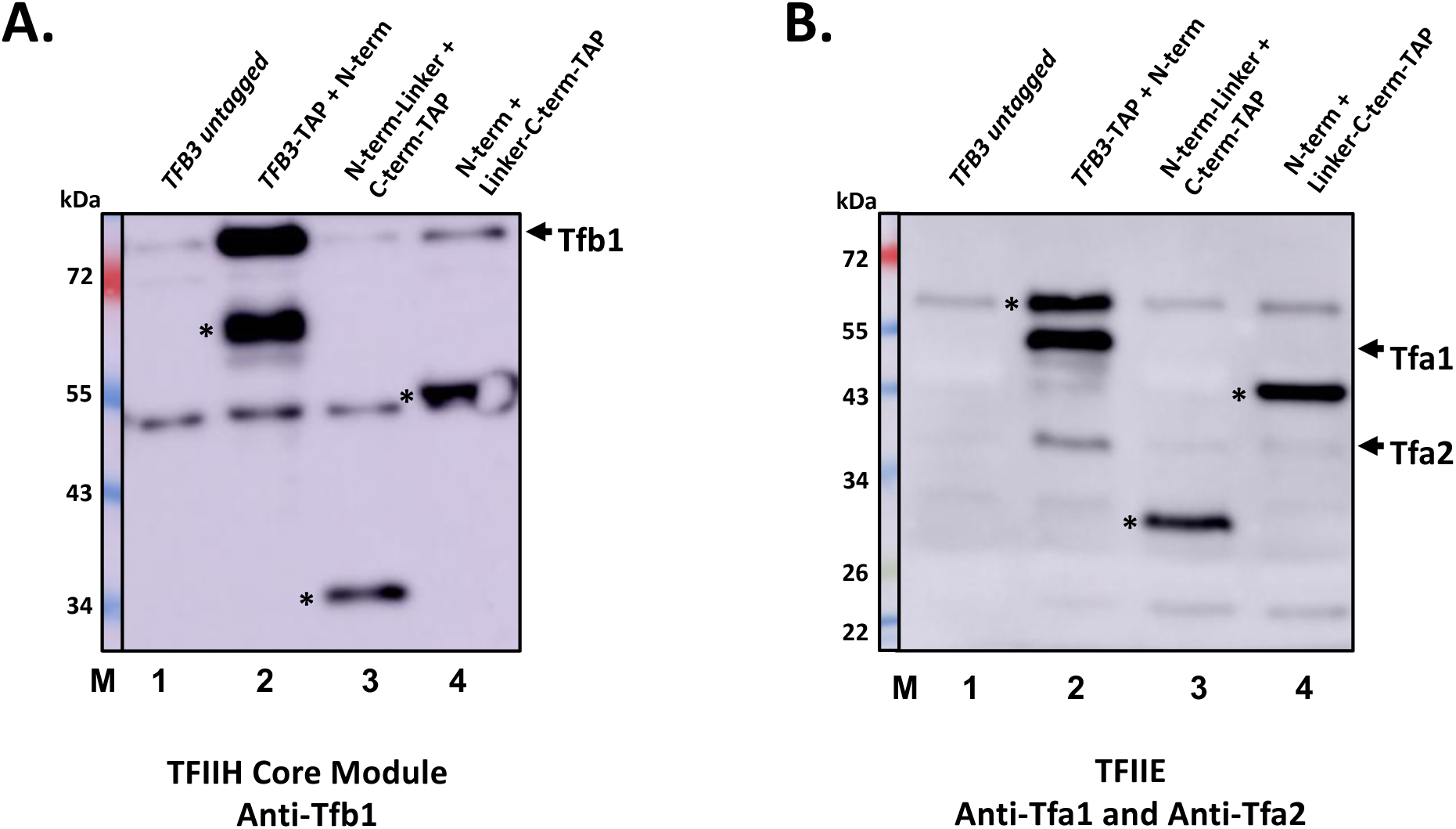
Splitting Tfb3 uncouples the TFIIH Kinase Module from the Core Module and the PIC. The same IgG-Agarose precipitates analyzed in Fig. 3 were also probed with the following antibodies against: **(A)** TFIIH Core Module subunit Tfb1 (note background binding of Tfb1 to beads as seen in lane 1, and non-specific band ∼50 kDa), and **(B)** TFIIE subunits Tfa1 and Tfa2 (note non-specific band ∼60 kDa), which serves as a marker for the RNApII PIC. In all panels, asterisks mark TAP-tagged Tfb3 derivatives reacting with secondary antibody due to the protein A component.

The basal transcription factor TFIIE directly contacts several Core TFIIH subunits, including Tfb3, within the RNApII PIC. Furthermore, TFIIE stimulates TFIIH enzymatic activities, including the kinase (Watanabe et al., 2003). To test how the TFIIE-TFIIH association was affected by the split Tfb3, the TAP-precipitated samples were also immunoblotted for the TFIIE subunits Tfa1 and Tfa2. Whereas full length Tfb3 efficiently pulls down both TFIIE subunits, the separated C-term or Linker-C-term derivatives do not **(Figure 4B)**. This result further indicates that the Kinase Module no longer associates with the TFIIH Core or the RNApII initiation complex upon splitting Tfb3 into two parts.

### Uncoupling the TFIIH modules disrupts proper kinase localization

Given the viability of the split Tfb3 strains, we wondered if the untethered Kinase Module was still delivered to promoters, perhaps via interaction with Mediator. Genome-wide chromatin immunoprecipitation (ChIP-Seq) was used to compare localization of the TFIIH modules in the wild-type and split Tfb3 strains. In cells expressing full-length Tfb3, the Core TFIIH subunit Tfb1 showed a strong peak near the transcription start site, reflecting promoter localization in the PIC (**Figure 5A and Figure 5 – figure supplement 1A**, red line). In all three split Tfb3 strains (yellow, blue, and green), the Tfb1 signal was reduced relative to the full-length, but still clearly promoter-localized. Perhaps reflecting reduced PIC formation, a similar reduction was seen for RNApII subunit Rpb1 (**Figure 5D and Figure 5 – figure supplement 1D**). Interestingly, the Tfb1 signal was closest to normal in the strain in which the Linker was fused to the N-terminal domain, despite the fact that this strain grew slower than strains with the Linker on the C-terminal domain or missing completely (**Figure 2C**). In contrast to Core TFIIH, the Kin28 ChIP signal, normally localized at promoters in wild-type Tfb3 strains, was completely absent in all the split Tfb3 strains (**Figure 5B** and **Figure 5 – figure supplement 1B**). Kin28 levels in extracts were below the limit of detection for our antibody, so we cannot rule out that the reduced ChIP signal is partly due to lower Kin28 levels in the split Tfb3 strains. However, the viability of the cells (**Figure 2**) and the Tfb3-TAP purifications (**Figure 3**) argue against a complete loss of Kin28. We conclude that, although Core TFIIH is still properly localized, the untethered Kinase Module is no longer efficiently targeted to promoters.

**Figure 5.**
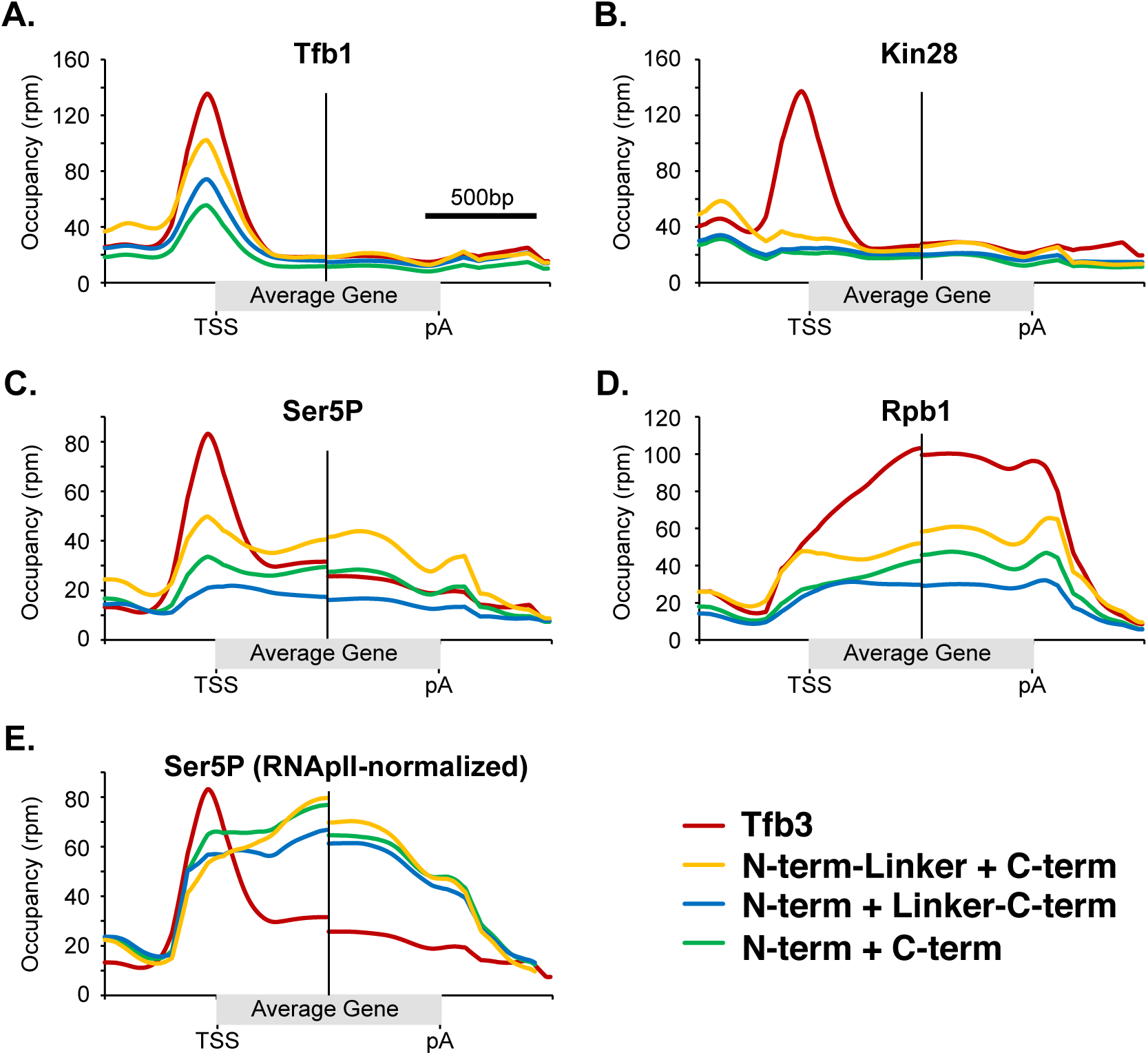
**Loss of proper Kinase Module and Ser5P promoter localization in split Tfb3 cells**. Crosslinked chromatin was prepared from strains expressing full length Tfb3 (YSB3788, red line), the separated N-term-Linker and C-term derivatives (YSB3731, yellow), the N-term and Linker-C-term derivatives (YSB3727, blue), or N-term and C-term with no Linker (YSB3725, green). Samples were immunoprecipitated and processed for sequencing of crosslinked DNA as described in Methods. Two biological replicates for each antibody were used to generate metagene profiles of **(A)** TFIIH Core subunit Tfb1, **(B)** TFIIH Kinase subunit Kin28, **(C)** Rpb1 CTD Ser5P, **(D)** RNApII subunit Rpb1, and **(E)** Ser5P normalized to total Rpb1. The analysis used only those nuclear genes with an average RNApII occupancy greater than 4 reads per million (rpm) and longer than 1 kb (n=95). Individual replicates are shown in Figure 5 **Supplement 1**. To account for the differing lengths of genes, each graph shows 1 kb centered on the transcription start site (TSS) to the left of the dividing line, and 1 kb centered on the polyadenylation site (pA) to the right.

### Untethered kinase module leads to aberrant Ser5P patterns

We expected that loss of Kinase Module recruitment would affect CTD Serine 5 phosphorylation (Ser5P) negatively, with possible indirect effects on Ser2P and overall elongation processivity. This prediction was first tested using immunoblotting of whole cell extracts. Cells lacking any Tfb3 C-term were strongly reduced for both Ser5P and Ser2P (**Figure 5 – figure supplement 2**, lanes 2 and 3). Similar to the growth phenotypes, adding back the C-term as a separate protein greatly improved CTD phosphorylation levels at both sites, albeit not to full wild-type levels (lanes 4 and 5).

ChIP-seq was used to determine the location of Ser5P, normalizing to spiked-in *S. pombe* chromatin. Surprisingly, the restored Ser5P was abnormally distributed in split Tfb3 cells. Total RNApII crosslinking (**Figure 5D and Figure 5 – figure supplements 1D and 3A, B**) was reduced in the mutants, but the levels remained consistent across the gene. This pattern argues against an elongation processivity defect in the split Tfb3 strains, and contrasts with the 5’ to 3’ drop seen upon complete depletion of Kin28 (Wong et al., 2014) or Bur2, the cyclin for the elongation kinase Bur1/Cdk9 (Keogh et al., 2003). Ser5P normally has a strong peak at promoters, and lower but significant levels throughout transcribed regions, as can be seen in both individual gene traces (**Figure 5 – figure supplement 3A, B**, promoters in green boxes) and metagene analysis (**Figure 5C** and **Figure 5 – figure supplement 5C**, red curves). In the split Tfb3 strains, the Ser5P promoter peak was missing, but levels of this modification remained high throughout the downstream transcribed region. Normalization of Ser5P to total RNApII indicated that downstream Ser5P was actually elevated in the mutants relative to wild-type (**Figure 5E**).

Quantitation of RNApII-normalized Ser5P at individual genes (**Figure 5 -figure supplement 1E, F**) showed that this redistribution of Ser5P occurred at virtually all genes. The Ser5P signals at promoters are reduced by up to 50%, particularly at strongly transcribed genes (i.e. those with highest RNApII occupancy), while the distribution for the mutant/WT ratio is centered around one for the weakest genes (**Figure 5 – figure supplement 1E**, left panels). In contrast, the normalized Ser5P signal in open reading frames (right panels) are nearly all increased by two-fold or more in the mutants. Despite the changes in CTD phosphorylation, the steady RNApII levels across genes in the mutants (**Figure 5D**) suggest no defect in elongation processivity. Using the total Rpb1 signals as a measure of transcription levels, it can be seen that transcription is clearly down overall in the split Tfb3 strains (**Figure 5 - figure supplement 1F**). Isolated analyses of genes that were longer than average, expressed at different levels, or that were more or less dependent on TFIID for initiation showed no differences from the total (**Figure 5 – figure supplement 4A-C**). Notably, seven genes showed an opposite effect from the general drop in transcription, with reproducibly increased Rpb1 signals in the mutants. All but one of these are known heat shock response genes (**Figure 5 – figure supplement 1G**), suggesting these slow growing cells are under stress.

ChIP-seq was also performed for Ser2P. While the overall signals were too low to get good quality data for metagene analysis, individual gene traces at strongly transcribed genes suggested that the pattern of Ser2P was relatively normal (**Figure 5 – figure supplement 3B**). In both WT and split Tfb3 strains, Ser2P was lowest near the promoter and gradually increased further downstream.

The surprising increase of Ser5P downstream of the promoter suggests that the Tfb3 linkage not only delivers the Kinase Module to the promoter, but may also inhibit its activity outside of the PIC. Transcription-associated phosphatases normally cause Ser5P to drop soon after RNApII enters the body of the gene (reviewed in (Buratowski, 2009; Corden, 2013; Eick & Geyer, 2013)), so the higher downstream Ser5P signal suggests untethered kinase module may be able to continually access the CTD during elongation. This could be due to inhibition of TFIIH Core-associated Kinase outside of the PIC, or suggest that the majority of TFIIH is sequestered at promoters and therefore unavailable at later stages of transcription.

### Split Tfb3 does not cause an NER defect

The TFIIH Core Module also functions as a key component of the NER machinery, and Tfb3 has been shown to be important for this function (Feaver et al., 2000). While many mutations in Core subunits therefore cause sensitivity to ultraviolet (UV) light, this is not the case for Kinase Module mutants. Recent structures (reviewed in (van Sluis et al., 2025; Yu et al., 2023)) suggest that the Kinase Module would block interactions between the Core Module and other NER factors. Therefore, TFIIH either enters into the NER complex as free Core Module, or the Kinase Module may dissociate soon after. We tested the response to UV light in strains where the split Tfb3 dissociates the Core and Kinase Modules (**Figure 6**). Two wild-type Tfb3 strains acted as controls for normal UV response, including one strain lacking the Tfb6 protein that competes with the Core for Ssl2 binding (Murakami et al., 2012), as this background was used for many experiments here (see **Supplemental Table S2**). None of the split Tfb3 configurations caused UV sensitivity, indicating that the separated N-term of Tfb3 suffices for the Core NER function. In fact, we noticed that strains with free N-term domain, where the linker is either absent or connected to the C-terminal domain, may even have slight resistance to UV light, consistent with a model in which the Kinase Module inhibits the TFIIH NER function. Any NER role for the kinase module must be functional even in the split Tfb3.

**Figure 6.**
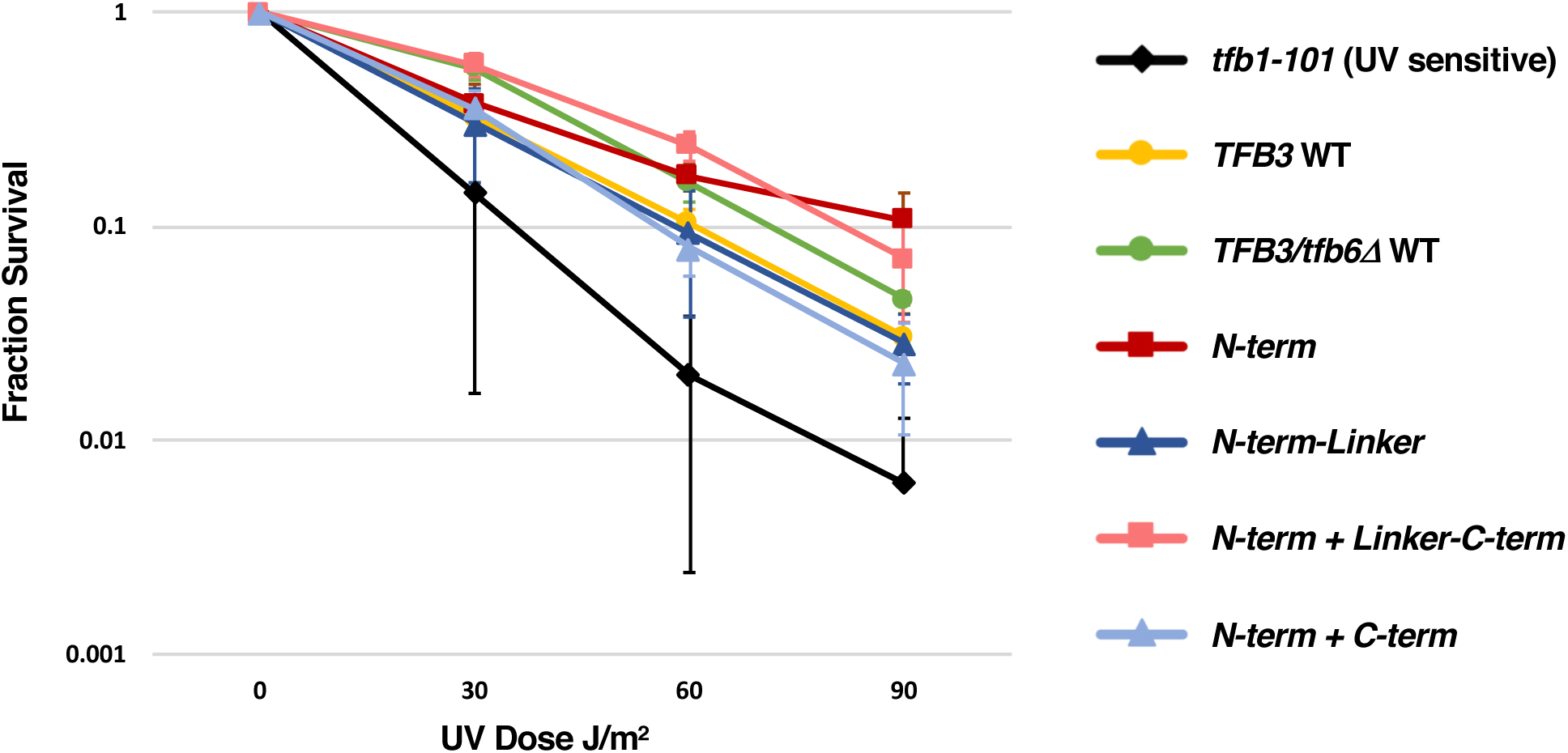
**Split Tfb3 strains are not hyper-sensitive to UV light**. Yeast strains carrying the indicated Tfb3 configurations were plated at various densities and exposed to the indicated dosages of UV light. Surviving colonies were counted after several days. The graph shows the averaged values from multiple replicates, and error bars represent standard deviation. Note that there are two strain backgrounds. The *tfb1-101* strain (YSB250, black) is a positive control for an NER-defective strain sensitive to UV light, and its isogenic WT control is YSB207 (yellow). The other five isogenic strains are in a *tfb6Δ* background (Warfield et al., 2016), lacking the Tfb6 protein that competes with the TFIIH for Ssl2 binding (Murakami et al., 2012): WT/tfb6Δ (YSB3788, green), N-term (YSB3724, red), N-term-Linker (YSB3729, blue), N-term + Linker-C-term (YSB3727, orange), and N-term-Linker + C-term (YSB3731, slate).

## DISCUSSION

We show here that the TFIIH Core and Kinase Modules can be uncoupled in vivo, leading to misregulated CTD phosphorylation (**Figure 2**). Although it was previously reported that both the N-term and C-term of Tfb3 are essential for viability in *S. cerevisiae* (Feaver et al., 2000; Warfield et al., 2016), we found that cells lacking the C-terminal domain are viable but extremely slow growing. Consistent with this finding, accurate transcription initiation can be reconstituted in vitro using a Tfb3 truncation consisting of only N-term residues 1-148 (Yang et al., 2022). Furthermore, Mouse Embryonic Fibroblasts and cardiomyocytes with a homozygous deletion of *MAT1* were shown to be viable but abnormal (Helenius et al., 2011; Sano et al., 2007). In these cells, Core TFIIH remains intact, but levels of the Kinase Module proteins CDK7 and Cyclin H are severely reduced. *MAT1* deletion leads to major defects in CTD phosphorylation and transcription, but the effects are partially compensated by reduced mRNA degradation rates. Therefore, our results splitting the Core and Kinase Modules in yeast are likely also relevant to mammalian TFIIH.

While fusion of other kinase modules to the Tfb3 linker did not substitute for the C-term (**Figure 1**), we found that adding back the Tfb3 C-terminal domain as a separate protein dramatically improved growth (**Figure 2**). As discussed below, this rescue apparently results from the restored CTD kinase activity of the untethered Tfb3 C-term/Kin28/Ccl1 complex (**Figure 3**, **Figure 3 – figure supplement 1, Figure 5 – figure supplement 2**). ChIP-Seq showed that Kin28 kinase was no longer preferentially recruited to promoters in the split Tfb3 strains, even though Core TFIIH still showed proper localization (**Figure 5 and Figure 5 – figure supplement 1**). Having uncoupled the Kinase Module from TFIIH, we expected CTD Ser5P to be severely reduced in the split Tfb3 strains. Indeed, Ser5P peaks at promoters were largely gone, but instead the modification was found throughout the downstream transcribed region at levels higher than normal **Figure 5**, **Figure 5 – figure supplements 1 and 3**).

Several models could explain the abnormal Ser5P pattern seen in strains with split Tfb3. One possibility is that a kinase other than Kin28 somehow substitutes for Ser5 phosphorylation, but without promoter-proximal targeting. However, several observations argue against a compensating kinase. First, any such kinase does not relieve the requirement for *KIN28* in supporting viability (**Figure 1C**). Second, reintroduction of the separated Tfb3 C-term markedly rescues CTD phosphorylation (**Figure 5 – figure supplement 2**), and it is unclear how a kinase other than Kin28 would respond in this manner. Finally, repeated attempts to introduce a chemically inhibitable Kin28 allele (Chun et al., 2019; Joo et al., 2019) into the split Tfb3 background were unsuccessful, and this apparent synthetic lethality further suggests that Kin28 supplies the essential Ser5 phosphorylation when the TFIIH modules are separated. We therefore consider the compensating kinase model unlikely.

A second possibility is that, under normal conditions, the majority of Kinase Module is tethered to the Core at promoters and thereby sequestered from phosphorylating downstream RNApII. Breaking the Tfb3 tether could free the Kinase Module to access the CTD throughout transcription. Arguing against this model, single molecule tracking in vivo reveals substantial fractions of freely diffusing Kin28 and core TFIIH (Nguyen et al., 2021). Furthermore, a significant amount of free CDK7/Cyclin H/MAT1 trimer exists in metazoans, providing CAK activity, and free trimer complex is also detected in yeast extracts (Keogh et al., 2002). On the other hand, the mammalian trimer complex appears to be largely cytoplasmic (Rimel et al., 2020). Therefore, a tethering model might be feasible if Kinase Module containing the Tfb3 C-term is more abundant in the nucleus than that with full-length Tfb3.

Both chromatin immunoprecipitation (Bataille et al., 2012; Komarnitsky et al., 2000; Mayer et al., 2010; Tietjen et al., 2010) and mass spectrometry (Suh et al., 2016) show that a low level of Ser5P persists during late elongation. The downstream Ser5P has been presumed to be residual phosphatase-resistant Ser5P acquired at the promoter. However, it is interesting to consider the possibility that free Kinase Module in the nucleus might prevent complete Ser5P dephosphorylation by providing additional de novo phosphorylation during late transcription.

This antagonism would be similar to CTD Ser2, where both Ctk1 kinase and Fcp1 phosphatase appear to function throughout elongation (Cho et al., 2001).

A third possible explanation for the increase in downstream Ser5P is that untethered Kinase Module can access the CTD during elongation in a way that intact TFIIH cannot. In addition to delivering the kinase to the initiation complex, the connected Core Module might limit CTD phosphorylation when TFIIH is outside of the PIC. Supporting this idea, we previously showed that CTD kinase activity of free trimer Kinase Module was less sensitive than TFIIH complex to mutations in the activating T-loop (Keogh et al., 2002). Also, Rimel et al. showed that the mammalian TFIIH core module significantly inhibits the activity of the mammalian CDK7/CycH/MAT1 complex, at least on substrates other than the CTD (Rimel et al., 2020). These results suggest the Core module has some restrictive effect on CTD kinase activity.

In addition to the Core modulating TFIIH Kinase activity, there appears to be communication in the other direction as well. Available TFIIH structures and our data suggest that the Tfb3 linker does more than simply tether the N- and C-terminal domains, as cells expressing a Linker-C-term fusion grew better than cells completely lacking the linker.

Comparison of the structures for free TFIIH complex, TFIIH incorporated into the PIC, and TFIIH as part of the NER machinery (reviewed in (van Sluis et al., 2025; Yu et al., 2023)) suggest a second possible linker function. The Core Module has a horseshoe shape, with the TFIIH ATPases, Ssl2/XPB and Rad3/XPD, at the two ends (see **Figure 7** schematic). The spacing between the ATPases changes substantially depending on context. In free TFIIH, the long alpha-helix in the Tfb3/MAT1 Linker spans from Tfb3 N-term, which contacts Rad3/XPD, to the other side of the horseshoe, where it directly contacts Ssl2/XPB (**Figure 7A**). Here, the central Linker domain may lock together the two TFIIH ATPases in a “closed” conformation. In the PIC, the Tfb3/MAT1 alpha-helix is disconnected from Ssl2/XPB, producing an “open” conformation with the Ssl2/XPB rotated outward, its active site bound to the double-stranded DNA downstream of RNApII (**Figure 7B**). TFIIH within the NER complex lacks the Kinase Module, and in this case the NER factors bring Rad3/XPD and Ssl2/XPB even closer together to facilitate the interaction of the damaged DNA with both ATPase active sites (van Sluis et al., 2025; Yu et al., 2023).

**Figure 7.**
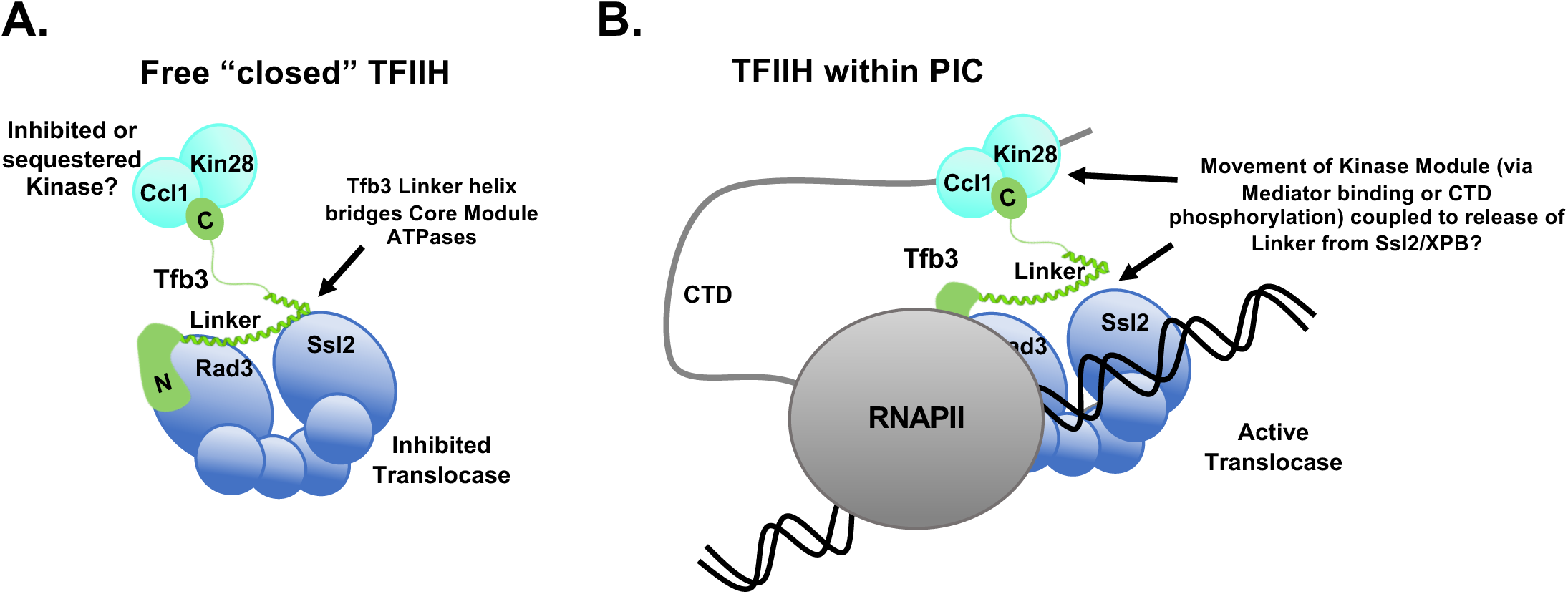
Model for linkage between the TFIIH Kinase and Core Module functions. **A.** In free TFIIH, the Tfb3 linker contacts the Ssl2/XPB subunit to create a “closed” conformation. In this configuration, the translocase, and possibly the kinase, may be inhibited. **B.** Upon TFIIH incorporation as the final component of the PIC, the Kinase Module engages with both Mediator (not pictured) and the CTD. These interactions may help release the Tfb3/MAT1 linker from Ssl2/XPB, allowing the translocase to access downstream DNA to promote ATP hydrolysis and promoter melting.

A report describing two PIC structures assembled in the context of a +1 nucleosome (Abril-Garrido et al., 2023) shows that when the nucleosome blocks Ssl2/XPB from downstream DNA, TFIIH is in the closed conformation, with the Tfb3/MAT1 Linker contacting both ATPases. Positioning the nucleosome further downstream allows the open TFIIH conformation, resulting in loss of the contact between the Tfb3/MAT1 Linker and Ssl2/XPB binding DNA. Notably, much of the Tfb3 long alpha-helix becomes disordered when disconnected from Ssl2/XPB. Although these two PIC structures lack Mediator, and it hasn’t been shown that the closed TFIIH PIC is a necessary precursor to the open TFIIH, it seems quite plausible that TFIIH enters the PIC in the closed conformation, transitioning to the open conformation upon DNA binding in order to proceed with DNA translocation and promoter melting.

Based on our findings, we speculate that the Tfb3 Linker provides a mechanism for temporal coupling of TFIIH kinase and promoter melting functions (**Figure 7B**). PICs with closed TFIIH are predicted to be inhibited for ATP hydrolysis by Ssl2/XPB (Abril-Garrido et al., 2023), and as described above, may also be partially inhibited for kinase activity (Keogh et al., 2002; Rimel et al., 2020). Movement of the Kinase Module during PIC assembly may reposition the Tfb3 Linker to release the inhibitory contact with Ssl2/XPB, freeing the translocase for promoter melting. This movement might be triggered by capture of the Kinase Module by Mediator, or possibly by initiation of CTD phosphorylation. Alternatively, DNA binding and translocation by Ssl2/XPB in the PIC might release the Linker contact to promote Kinase Module activity. CTD phosphorylation triggers release of Mediator (Sogaard & Svejstrup, 2007), and in vivo experiments with TFIIH kinase inhibition suggest this transition occurs very rapidly after PIC formation (Jeronimo & Robert, 2014; Wong et al., 2014). Notably, release of the Tfb3 Linker contact also results in the long alpha-helix becoming disordered (Abril-Garrido et al., 2023), which could allow the kinase access to a far larger radius of area. This flexibility could help the kinase reach both proximal and distal repeats within the CTD, which can theoretically extend quite far from the RNApII body. Supporting the idea that the Kinase Module is mobile, PICs formed in vitro without Mediator show the Kinase Module close to the Rpb4/Rpb7 stalk of RNApII (Abril-Garrido et al., 2023; Chen et al., 2021), far from its position seen when interacting with Mediator.

One species-specific feature of yeast PICs is that they can access transcription start sites (TSSs) far further downstream than metazoan PICs. Enzymatic studies show that for yeast TFIIH, but not its mammalian homolog, the Kinase Module greatly stimulates the processivity of the Ssl2 translocase activity necessary for downstream TSS scanning (Tomko et al., 2021), likely via the interaction of Tfb3 with RNApII. Yeast transcription reconstituted in vitro with TFIIH lacking Tfb3 and the Kinase Module causes a dramatic upstream shift in transcription start sites, with the TATA-TSS spacing now resembling that of metazoans (Murakami et al., 2015). This effect can be reversed by adding back only the Tfb3 N-term (Yang et al., 2022), arguing against a model where the contact between Tfb3 linker and Ssl2 is necessary for downstream TSS usage. Nevertheless, there is clearly some communication between the Kinase and Core Modules, and Tfb3 is likely to be a key link.

### Evolutionary considerations

We speculate that cells with a complete deletion of the Tfb3 Linker may resemble an evolutionarily ancient state where TFIIH Kinase and Core existed as completely distinct enzymes. Initially, the primary role of TFIIH Core was probably NER, and the Kinase Module precursor likely functioned to phosphorylate CDKs or other substrates. Each module would have then been independently co-opted for RNApII transcription. Supporting this hypothesis, TFIIH from the early diverged eukaryote *Trypanosome brucei* appears to consist of only the Core Module, as no Tfb3 or other Kinase Module homologs are detected in the purified enzyme complex or by sequence similarity search (Lee et al., 2010; Lee et al., 2009). The CTD of *T. brucei* Rpb1 is quite diverged, without the typical repeat sequence found in most eukaryotes (Badjatia et al., 2013; Das & Bellofatto, 2009). Similarly, protein sequence searches find no unambiguous TFIIH kinase/cyclin homologs among the cyclin-dependent kinases in *T. brucei* or *Giardia lamblia* (Guo & Stiller, 2004, 2005). In *Plasmodium falciparum*, which also lacks a consensus CTD, a putative Tfb3/MAT1 homolog has been described (Callebaut et al., 2005; Chen et al., 2006). However, this protein has not yet been shown to associate with TFIIH and there is still some uncertainty whether an associated kinase is homologous to Kin28/CDK7 or another CDK.

Overall, it appears likely the Tfb3 Linker evolved very early in the eukaryotic lineage, perhaps to coordinate the timing of CTD phosphorylation and promoter melting. In the split Tfb3 strains, fusion of Linker to the C-term domain may improve growth relative to the complete Linker deletion (**Figure 2C**) by partially restoring kinase-translocase coupling. Our ChIP-Seq results suggest an ancestral, free CTD kinase may have acted throughout elongation. Tethering the kinase to TFIIH would have targeted high levels of CTD phosphorylation during early transcription, establishing the first step of the sequential CTD cycle as we know it today, where distinct phosphorylation patterns mark the progression from early through late transcription.

## METHODS

| **Key Resources Table** |  |  |  |  |
| --- | --- | --- | --- | --- |
| **Reagent type (species) or resource** | **Designation** | **Source or reference** | **Identifiers** | **Additional information** |
| antibody | anti-Ccl1 (rabbit polyclonal) | Keogh et al., 2002 |  | (1:1000) |
| antibody | anti-Tfb3 (rabbit polyclonal) | Keogh et al., 2002 |  | (1:1000) |
| antibody | anti-Kin28 (rabbit polyclonal) | Covance | Cat# PRB-250C-500 | (1:1000) |
| antibody | anti-Tfb1 (rabbit polyclonal) | Matsui et al., 1995 |  | (1:1000) |
| antibody | anti-Tfa1 (rabbit polyclonal) | Kuldell & Buratowski 1997 |  | (1:1000) |
| antibody | anti-Tfa2 (rabbit polyclonal) | Kuldell & Buratowski 1997 |  | (1:1000) |
| antibody | anti-RPB1-phospho-CTD Ser2P (rat monoclonal 3E10) | Thermo Fisher/Chapman et al., 2007 | Cat# 14-9802-82 | (1:1000) |
| antibody | anti-RPB1-phospho-CTD Ser5 (Rat monoclonal 3E8) | Sigma/Chapman et al., 2007 | Cat# 04-1572-1 | (1:1000) |
| antibody | anti-RPB1-CTD (mouse monoclonal 8WG16) | Biolegend | Cat# 664906 | (1:1000) |
| antibody | anti-Rpb3 (Mouse monoclonal 1Y26) | Biolegend | Cat# W0012 | (1:1000) |
| antibody | anti-rabbit IgG peroxidase (goat polyclonal) | Sigma | Cat#: A0545-1ML | Use recommended dilution |
| antibody | anti-rat IgG peroxidase (goat polyclonal) | Sigma | Cat# A9037-1ML | Use recommended dilution |
| antibody | anti-mouse IgG peroxidase (goat polyclonal) | Sigma | Cat# A2304-1ML | Use recommended dilution |
| commercial assay or kit | Ni-NTA agarose | Gold Biotechnology | Cat# H-350-100 |  |
| commercial assay or kit | AminoLink Plus Coupling Resin | Thermo Fisher | Cat# PI20505 |  |
| commercial assay or kit | SuperSignal Plus Pico Kit | Thermo Fisher | Cat# PI34580 |  |
| commercial assay or kit | Protein G Dynabeads | Thermo Fisher | Cat. # 10004D |  |
| commercial assay<br>or kit | Pan Mouse IgG<br>Dynabeads | Thermo Fisher | Cat#<br>11042 |  |
| commercial assay<br>or kit | IgG Sepharose | Cytiva | Cat#<br>17096901 |  |
| commercial assay<br>or kit | KAPA Pure<br>Beads | Roche | Cat#<br>07983280001 |  |
| commercial assay<br>or kit | KOD Hot Start<br>DNA<br>Polymerase | Millipore | Cat#<br>71086-3 |  |
| commercial assay<br>or kit | NEBNext® Libr<br>ary Quant Kit<br>for Illumina® | New England<br>Biolabs | E7630 |  |

### Plasmid Construction and Yeast Strains

Plasmids used in this study, and their construction, are listed in **Supplemental Table S1**. Yeast strains used in this study are listed in **Supplemental Table S2**. All are available from the corresponding author upon request.

### Antibodies and Immunoblotting

The following primary antisera were used: anti-Ccl1 (Keogh et al., 2002), anti-Kin28 (Covance, PRB-250C-500), anti-Tfb1 (Matsui et al., 1995), anti-Tfa1 and anti-Tfa2 (Kuldell & Buratowski, 1997), anti-Ser2P (3E10), anti-Ser5P (3E8) (gift from Dirk Eick (Chapman et al., 2007)), anti-CTD (8WG16), anti-Rpb3 monoclonal 1Y26 (BioLegend). For immunoblotting, primary antibodies were used at a 1:1000 dilution. Secondary antibodies coupled to horse radish peroxidase (goat anti-rabbit, goat anti-rat, and goat anti-mouse) were purchased from Sigma and used at the recommended dilution.

Anti-Tfb3 antibodies (Keogh et al., 2002) were affinity purified from sera to reduce cross-reacting signals. Briefly, recombinant Tfb3 protein was expressed from plasmid pSBEThis7-Tfb3 in *E. coli* BL21(DE3). Cells were grown at 37°C to an OD_600_ of 0.4-0.6 in 4 L of selective media. Expression was induced with 0.2 mM IPTG and cultures were incubated shaking at 30°C for 4 hrs. Cells were pelleted and resuspended in 100 mL lysis buffer (1X PBS, 1% SDS, pH 7.4). After sonication, cell debris was pelleted at 23,970 g for 20 min. The supernatant was filtered and then incubated with nutation at 4°C for 1 hr with Ni-NTA agarose (Gold Biotechnology #H-350-100) that was pre-equilibrated with lysis buffer. Lysate and beads were drained through a column and washed extensively with wash buffer (1X Phosphate Buffered Saline (PBS), 0.1% Sarkosyl, 10 mM imidazole, pH 7.4). Bound protein was serially eluted with elution buffer (1X PBS, 0.1% Sarkosyl, pH 7.4) supplemented with increasing imidazole concentrations (1 mL each): 50 mM, 100 mM, 150 mM (2X), 200 mM, 300 mM.

Fractions with the highest Tfb3 yield were pooled. Roughly 2 mg of purified Tfb3 was covalently linked to 1 ml AminoLink® Plus Coupling Resin (Thermo Scientific #PI20505) following the manufacturer’s protocol for crosslinking at pH 10 with cyanoborohydride solution. Specific anti-Tfb3 antibodies were affinity purified from crude antiserum as described (Harlow & Lane, 1988). Tfb3-coupled beads were serially washed with 10 bed volumes each of 10 mM Tris pH 7.5, 100 mM glycine pH 2.5, 10 mM Tris pH 8.8, and then 100 mM triethylamine pH 11.5.

Finally, beads were poured into a small column and washed with 10 mM Tris pH 7.5 until pH reached 7.5. Antiserum was diluted 1:10 with 10 mM Tris pH 7.5 and passed through the column three times. The column was then washed with 20 bed volumes each of 10 mM Tris pH 7.5 and 500 mM NaCl, and then 10 mM Tris pH 7.5. Bound antibodies were serially eluted with small volumes of high (100 mM triethylamine pH 11.5) and low (100 mM glycine pH 2.5) pH, followed by rapid neutralization. Peak fractions were pooled and concentrated using Amicon Ultra 0.5 10K Concentrator, with PBS as the replacement buffer.

### Yeast whole cell extracts

Two types of extracts were used for gel electrophoresis and immunoblotting, with similar results. The first is an alkaline lysis method. 1.5 mL of late log phase liquid culture was pelleted at 4,600 g for 90 sec at room temperature. Media was removed, and the pellet was resuspended in 0.1 M NaOH and incubated at room temperature for 5 min. Cells were pelleted at 9,300 g for 90 sec at room temperature, supernatant was removed, and pellet was resuspended in 60 μL lysis buffer (0.06 M Tris-HCl pH 6.8, 5% glycerol, 2% SDS, 4% β-mercaptoethanol, 0.0025% bromophenol blue) (Kushnirov, 2000). Samples were heated at 95°C for 5 min, pelleted at 9,300 g for 1:30 min at room temperature, and supernatant was collected to run on an SDS-PAGE gel.

For some gels, a non-denaturing whole cell extract was used. First, 1.0-1.5 mL of liquid culture was pelleted (all centrifugations were at 2,300 g) at 4°C for 2 min and washed with 1 mL cold distilled water. Pellets were resuspended in 100 μL cold lysis buffer (50 mM HEPES pH 7.6, 1 mM EDTA, 300 mM NaCl, 20% glycerol, 1 mM DTT, 1 μg/mL leupeptin, 1 μg/mL aprotinin, 1 μg/mL pepstatin A, and 1 μg/mL antipain). The resuspension was mixed with 100 μL 425-600 μm glass beads (Sigma, #G8772-1KG) and vortexed at 4°C for 10 min. Extract was centrifuged for 10 min at 4°C and supernatant collected. Total protein concentration of extracts was measured using NanoDrop 2000 Spectrophotometer (ThermoScientific).

### TAP purification and co-precipitation assay

TAP-tagged Tfb3 derivatives were used for assaying Kinase Module association and in vitro kinase assays. Yeast strains were grown in 200 mL of selective media at 30°C to an OD_600_ of ∼2. Cells were collected by centrifugation, flash frozen in liquid nitrogen, and lysed using a SPEX SamplePrep Freezer Mill 6875D with small chambers. Pulverized cells were resuspended by rotation at 4°C in 1 mL of lysis buffer (20 mM HEPES pH 7.6, 10% glycerol, 200 mM KoAc, 1 mM EDTA, 1 mM DTT, 1 mM NaF, 1 mM Na3VO4, 1 μg/mL leupeptin, 1 μg/mL aprotinin, 1 μg/mL pepstatin A, 1 μg/mL antipain, 1 mM PMSF) until homogeneous. Cell debris was pelleted in a benchtop microfuge at 4,500 g for 25 min, and supernatant was collected and further cleared in a microfuge at maximum speed for 10 min. All centrifugations were done at 4°C. The clear layer of lysate was transferred to a new tube, 10 μL of IgG Sepharose (GE Healthcare, #17096901) pre-equilibrated with lysis buffer was added, and the suspension was rotated at 4°C for 1 hr. After binding, IgG beads were gently pelleted, supernatant was removed, and beads were washed 3 times with lysis buffer.

For analysis of bound proteins, 50 μL of 2X SDS-PAGE sample buffer (0.1% bromophenol blue, 0.2 M DTT, 20% glycerol, 4% SDS, 0.1 M Tris-Cl pH 6.8) were added to the beads and the sample was heated at 100°C for 5 min. The beads were then pelleted by centrifugation, supernatant was collected, and samples were analyzed by immunoblotting. Input, flowthrough, and final samples from the co-immunoprecipitation protocol were run on 10% polyacrylamide SDS-PAGE gels. Proteins were transferred to polyvinylidene difluoride (PVDF) membranes.

Membranes were blocked with 5% powdered milk in Tris Buffered Saline with 0.1% Tween and then probed with the appropriate primary antibody. Blocking and antibody incubations were done for either 1 hr at room temperature or overnight at 4°C. Following several washes, the blots were incubated with secondary antibody at a 1:10,000 dilution for 1 hr. Reacting bands were detected by chemiluminescent detection using the Thermo SuperSignal Plus Pico kit according to the manufacturer’s instructions.

### In vitro CTD phosphorylation assay

IgG beads were used to immobilize the Tfb3-TAP derivatives as described above. After normalizing for the amount of Kin28, beads were incubated with 100 ng GST-CTD in 1x kinase reaction buffer (20 mM HEPES pH 7.6, 7.5 mM MgOAc, 2% Glycerol, 100 mM KOAc, 2 mM DTT, and 25 μM ATP). Samples were taken at 15, 30, and 60 min, and phosphorylation was assayed by immunoblotting with anti-Ser5P (3E8) antibody.

### ChIP-seq

ChIP experiments were performed from two independent biological replicates as previously described (Jeronimo et al., 2021). Yeast cultures were grown in 50 mL of selective medium at 30°C to an OD_600_ of 0.7-0.9 before crosslinking with 1% formaldehyde (Fisher Scientific, BP531-500) at room temperature for 30 min and quenched with 125 mM glycine.

Crosslinked cells were collected by centrifugation and washed twice with 1X TBS (20 mM Tris-HCl pH 7.5, 150 mM NaCl). Cell pellets were then resuspended in 700 μL Lysis buffer (50 mM HEPES-KOH pH 7.5, 140 mM NaCl, 1 mM EDTA, 1% Triton X-100, 0.1% Na-deoxycholate and protease inhibitor cocktail (1 mM PMSF, 1 mM benzamidine, 10 μg/mL aprotinin, 1 μg/mL leupeptin, 1 μg/mL pepstatin A)). About the same number of OD_600_ units was used for all samples. A fixed amount of crosslinked *Schizosaccharomyces pombe* cells (a gift from Jason Tanny), at 10% the amount of *S. cerevisiae* cells, was added for normalization as previously described (Jeronimo et al., 2019). Cells were lysed by bead beating and the lysate was sonicated with a Model 100 Sonic dismembrator equipped with a microprobe (Fisher Scientific), 4 x 20 sec at output 7 Watts, with a 1 min break between sonication cycles. Soluble fragmented chromatin was recovered by centrifugation. 600 μL of the chromatin sample was taken per immunoprecipitation (IP) and 6 μL (1%) was saved as an Input sample. The following amounts of antibody per IP were used: anti-RNApII Rpb1 8WG16 (2 μg), anti-Ser5P 3E8 (5 μL), anti-Tfb1 (4 μL) and anti-Kin28 (4 μL). All antibodies have been validated for ChIP. Anti-Ser5P 3E8, anti-Tfb1, and anti-Kin28 antibodies were coupled to Dynabeads coated with Protein G (Thermo Fisher Scientific, 10004D) and anti-RNApII Rpb1 8WG16 antibody was coupled to Dynabeads coated with Pan Mouse IgG antibodies (Thermo Fisher Scientific, 11042). 50 μL of the appropriate Dynabeads pre-coupled with the indicated antibody were added to the chromatin sample and incubated overnight at 4°C. Beads were washed twice with Lysis buffer, twice with Lysis buffer 500 (Lysis buffer + 360 mM NaCl), twice with Wash buffer (10 mM Tris-HCl pH 8.0, 250 mM LiCl, 0.5% NP40, 0.5% Na-deoxycholate, 1 mM EDTA) and once with TE (10 mM Tris-HCl pH 8.0, 1 mM EDTA). Input and immunoprecipitated chromatin were eluted and reverse-crosslinked with 50 μL TE/SDS (TE + 1% SDS) by incubating overnight at 65°C. Samples were treated with RNase A (345 μL TE, 3 μL 10 mg/mL RNAse A (Sigma-Aldrich, R6513), 2 μL 20 mg/mL Glycogen (Roche, 10901393001)) at 37°C for 2 hr and subsequently subjected to Proteinase K (15 μL 10% SDS, 7.5 μL 20 mg/mL Proteinase K (Thermo Fisher Scientific, 25530049)) digestion at 37°C for 2 hr. Samples were extracted twice with phenol/chloroform/isoamyl alcohol (25:24:1), followed by precipitation with 200 mM NaCl and 70% ethanol. Precipitated DNA was resuspended in 50 μL of TE before being used in ChIP-seq libraries.

All ChIP and Input samples were subjected to a 1.7X cleanup using KAPA Pure Beads (Roche, 07983280001) according to the manufacturer’s instructions. Samples were eluted with 40 μL of Elution buffer (10 mM Tris pH 8.0). DNA concentration of ChIP and Input samples was determined by qPCR using a standard curve made with a fragment size control sample. 1-10 ng of ChIP and Input DNA were used for ChIP-seq library preparation as follows. The ends of DNA were repaired by incubating in 70 μL of 1X NEBuffer 2 containing 0.6 units of T4 DNA polymerase (NEB, M0203S), 2 units of T4 polynucleotide kinase (NEB, M0201S), 0.09 nM dNTPs and 0.045 μg/μL of BSA at 12°C for 30 min. Repaired DNA was then subjected to a 1.7X cleanup using KAPA Pure Beads before dA tailing as follows. Beads containing the repaired DNA were resuspended in 50 μL of 1X NEBuffer 2 containing 0.1 mM dATP and 25 units of Klenow Fragment (3’→5’ exo-) (NEB, M0212M), and incubated at 37°C for 30 min. After a 1.8X cleanup with KAPA Pure Beads, the A-tailed DNA was ligated to index adapters (Illumina Truseq DNA UD Indexes (20023784)) as follows. The beads were resuspended in 45 μL of 1X Ligase buffer containing 8 nM of adapter and 2.5 units of T4 DNA ligase (ThermoFisher, 15224041) and incubated at room temperature for 60 min. The ligated DNA was then subjected to a 1X cleanup with KAPA Pure Beads, followed by a double size selection (0.52X-1X) leading to fragments in the 200-600 bp range. Libraries were PCR-amplified with 12-13 cycles using KOD Hot Start DNA polymerase (Millipore, 71086-3) and cleaned up using 1X KAPA Pure Beads.

Libraries were qualified on Agilent 2100 Bioanalyzer using High Sensitivity DNA Kit and quantified by qPCR using NEBNext Library Quant Kit for Illumina (NEB, E7630). Equal molarity of each library was pooled and subjected to sequencing on Illumina NovaSeq 6000 platform at the McGill University and Génome Québec Innovation Centre to generate 100 bp paired-end reads.

### ChIP-seq Data analysis

ChIP-seq data was analyzed as previously described (Jeronimo et al., 2021). Adapter sequences were removed from paired-end reads for each biological replicate using Trimmomatic, version 0.36 (Bolger et al., 2014) with parameters “ILLUMINACLIP:adapters.fa:2:30:10: TRAILING:3 MINLEN:25“. Paired-end reads for each biological replicate were independently aligned to the *S. cerevisiae* (UCSC sacCer3) and *S. pombe* (Downloaded from https://www.pombase.org/ on December 7th 2022) reference genomes using the short read aligner Bowtie 2 (version 2.3.4.3) (Langmead et al., 2009). Only reads mapped in proper pairs and primary alignments were kept in aligned files using Samtools, version 1.9 (Li et al., 2009) with parameters “-F 2048 -F 256 -f 2”. Duplicate reads were removed from aligned files using Samtools “fixmate” and “markdup -r”. Coverage for each base pair of the *S. cerevisiae* genome was computed using genomeCoverageBed from BEDTools, version 2.27.1 (Quinlan & Hall, 2010) and normalized using *S. pombe* reads as follows. The read density at each position of the *S. cerevisiae* genome was multiplied by a normalization factor N defined as:

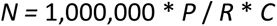

where:

1,000,000 is an arbitrarily chosen number used for convenience;

*R* is the total number of reads in the IP sample that mapped to the *S. pombe* genome; *C* is the total number of reads in the Input sample that mapped to the *S. cerevisiae* genome;

*P* is the total number of reads in the Input sample that mapped to the *S. pombe* genome.

The deduplicated BAM files of replicates were merged using Samtools “merge”.

Correlation between replicates were generated using deepTools, version 3.5.1 (Ramirez et al., 2016) with parameters “multiBigwigSummary bins --binSize 100” and “plotCorrelation --corMethod pearson --whatToPlot heatmap --colorMap PiYG --plotNumbers”.

The genome coverage files were converted to the bigWig format using the utilities from UCSC Genome Browser (Casper et al., 2018).

Statistics regarding the reads (including the number of reads, mapped reads, the correlation between replicates, etc.) are shown in **Supplemental Table S3**.

The RNApII-adjusted Ser5P coverage file (used in **Figure 5E** and **Figure 5 – figure supplement 1E**) was generated as follows, starting from *S. pombe*-normalized coverage files. For each genomic position, the Ser5P occupancy value of a given Tfb3 mutant was multiplied by the RNApII occupancy value in WT and divided by the RNApII occupancy value in the mutant.

Aggregate profiles (metagene analyses) were generated using the Versatile Aggregate Profiler (VAP) (Brunelle et al., 2015; Coulombe et al., 2014). The aggregate profiles were generated by aligning data on the transcription start site (TSS) and the polyadenylation (pA) signal of all verified genes that have a signal of RNApII >4 and are longer than 1 kb (n=95). The data were averaged over 10 bp bins. TSS and pA site coordinates were from Pelechano et al. (Pelechano et al., 2013).

In **Figure 5 – figure supplement 1E**, the values for “promoters” are the average signal over a 200 bp window centered on the TSS whereas the values for “ORFs” are the average of the signal between position +200 and the pA site. The region between +100 and +200 was excluded to avoid signal contamination from promoters for ORFs and vice versa. In **Figure 5 – figure supplement 4**, gene classifications are from (Donczew et al., 2020).

All raw and processed ChIP-seq data generated in this study have been deposited in the NCBI Gene Expression Omnibus (GEO; http://www.ncbi.nlm.nih.gov/geo/) under accession number GSE241436.

### UV sensitivity assay

Yeast strains were grown in liquid media selective for the *TFB3* plasmids, and overnight cultures were diluted to an OD_600_ = 0.005 and 0.0005 for plating. For each test plate, 100 μL of diluted cells were spread on selective media. Plated cells were then exposed to one of four doses of UV radiation: 0, 30, 60, or 90 J/m^2^. UV intensity was calibrated using a UVX Radiometer UVP and dosage calculated by varying time of exposure to a short-wave UV transilluminator. After irradiation, plates were grown at 30°C for 2 days in the dark to avoid photoreactivation repair, and then at room temperature for 5 days. Individual colonies were counted and fraction survivals were calculated and plotted.

## Article and Author Information

### Author details Gabriela Giordano

Department of Biological Chemistry and Molecular Pharmacology, Harvard Medical School, Boston, MA, USA

**Contribution**:; Investigation; Writing – original draft preparation; Visualisation

**Competing interests**: No competing interests declared.

### Robin Buratowski

Department of Biological Chemistry and Molecular Pharmacology, Harvard Medical School, Boston, MA, USA

**Contribution**: Investigation; Project administration

**Competing interests**: No competing interests declared.

### Célia Jeronimo

Institut de recherches cliniques de Montréal (IRCM), Montréal, Québec, Canada

**Contribution**: Formal analysis; Data curation; Investigation; Writing – review & editing; Visualisation

**Competing interests**: No competing interests declared.

### Christian Poitras

Institut de recherches cliniques de Montréal (IRCM), Montréal, Québec, Canada

**Contribution**: Formal analysis

**Competing interests**: No competing interests declared.

### François Robert

Institut de recherches cliniques de Montréal (IRCM), Montréal, Québec, Canada Département de Médecine, Université de Montréal, Montréal, Québec, Canada

Division of Experimental Medicine, Medicine, McGill University, Montréal, Québec, Canada

**Contribution**: Conceptualization; Data curation; Formal analysis; Writing – review & editing; Supervision; Funding acquisition

**Competing interests**: No competing interests declared.

### Stephen Buratowski

Department of Biological Chemistry and Molecular Pharmacology, Harvard Medical School, Boston, MA, USA **Contribution**: Conceptualization; Investigation; Writing – original draft preparation; Writing – review & editing; Visualisation; Supervision; Project administration; Funding acquisition

**Competing interests**: No competing interests declared.

## Funding

National Institutes of Health (R01GM046498)

Stephen Buratowski

### National Institutes of Health (R01GM056663)

Stephen Buratowski

### Canadian Institutes of Health Research (PJT-162334)

François Robert

The funders had no role in study design, data collection and interpretation, or the decision to submit the work for publication.

## ACKNOWLEDGEMENTS

We are grateful to Steve Hahn (FHCRC, Seattle) and John Feaver (Stanford) for providing several Tfb3 strains and plasmids, and to Jason Tanny (McGill Univ., Montreal) for *S. pombe* cells.

---

**Figure 1 – figure supplement 1.**
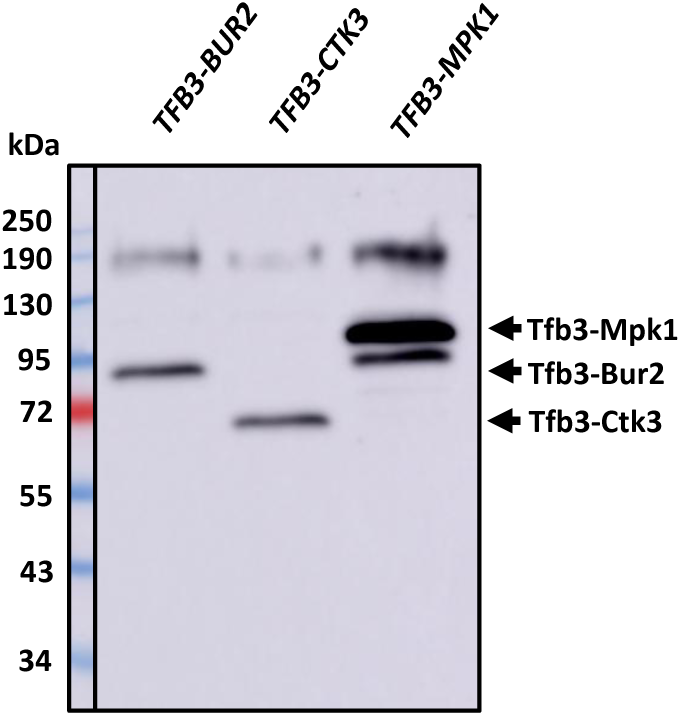
Immunoblot of Tfb3 fusions from Fig. 1C. Whole extracts were made from the post-FOA strains constructed by shuffling pRS425-*TFB3-BUR2*, pRS425-*TFB3-CTK3*, or pRS425-*TFB3-MPK1* into *tfb3Δ* strain SHY907/YF2456 (Warfield et al., 2016). Extracts were resolved by SDS-PAGE, blotted to a membrane, and probed with anti-Tfb3 antiserum. Left-most lane shows markers imaged with visible light, and the rest of the blot shows chemiluminescence detection.

**Figure 3 – figure supplement 1.**
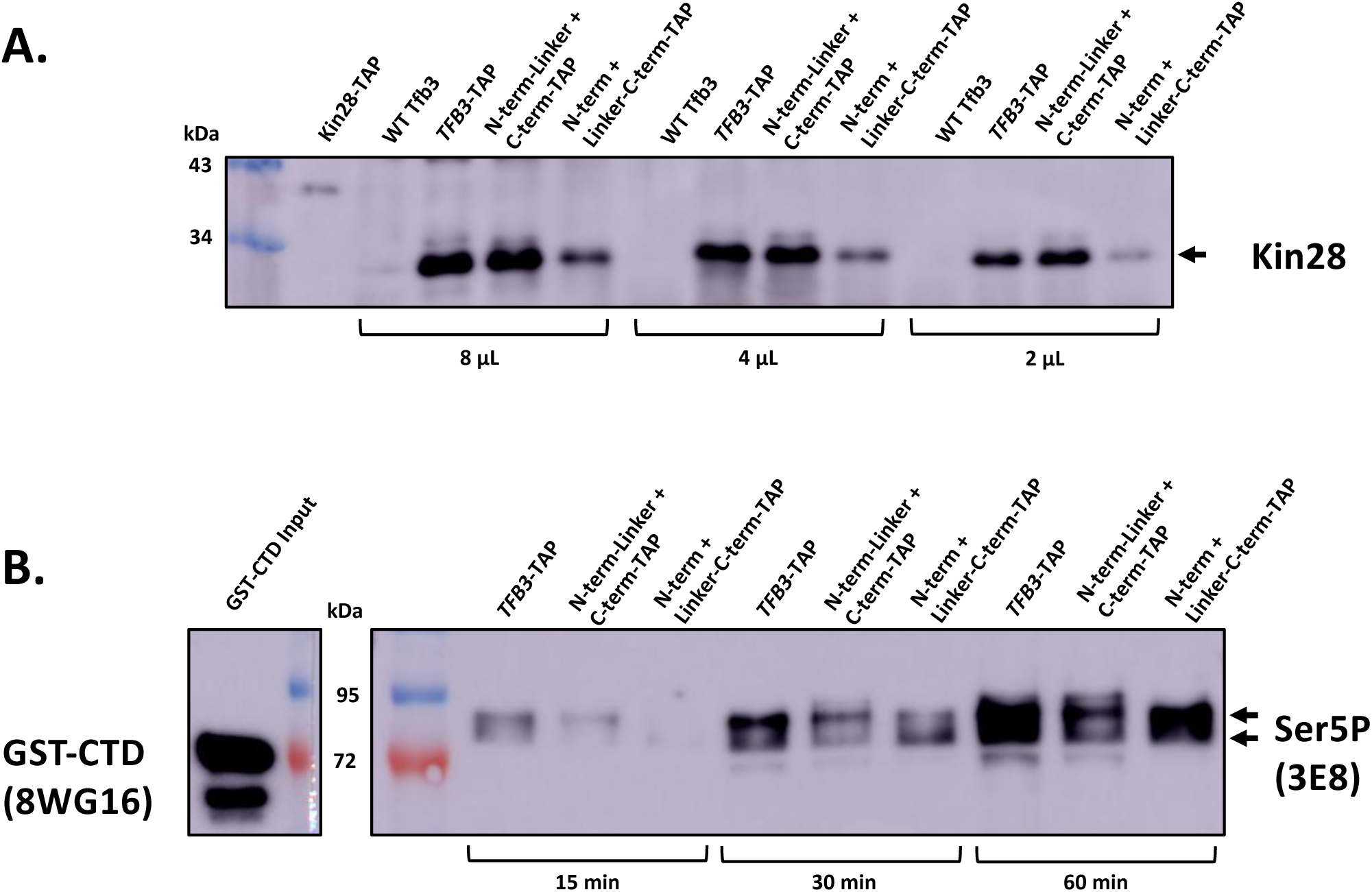
Tethered and untethered Kinase Module have similar CTD kinase specific activity in vitro. **(A)** Kinase Module was isolated from YSB3707 (TFB3-TAP), YSB3732 (N-term-Linker + C-term-TAP), and YSB3728 (N-term + Linker-C-term-TAP) using the TAP tag. A parallel isolation from the untagged strain YSB3788 (WT TFB3) showed that binding was dependent on the TAP tag. The second lane shows a Kin28-TAP precipitate (not done in parallel) as size marker. Relative levels of the Kin28 subunit on beads were determined by immunoblotting 8uL, 4uL, and 2uL of beads. **(B)** In vitro CTD phosphorylation assay. After roughly normalizing bead volumes for amount of bound Kin28, kinase reactions were performed with GST-CTD. Samples were taken at the indicated time points and tested for CTD Ser5 phosphorylation using monoclonal antibody 3E8. Note the phosphorylated CTD as a series of bands, with slower mobility believed to represent denser phosphorylation. To mark the position of unphosphorylated GST-CTD, a negative control reaction lane from the blot was probed with 8WG16 (left panel).

**Figure 5 – figure supplement 1.**
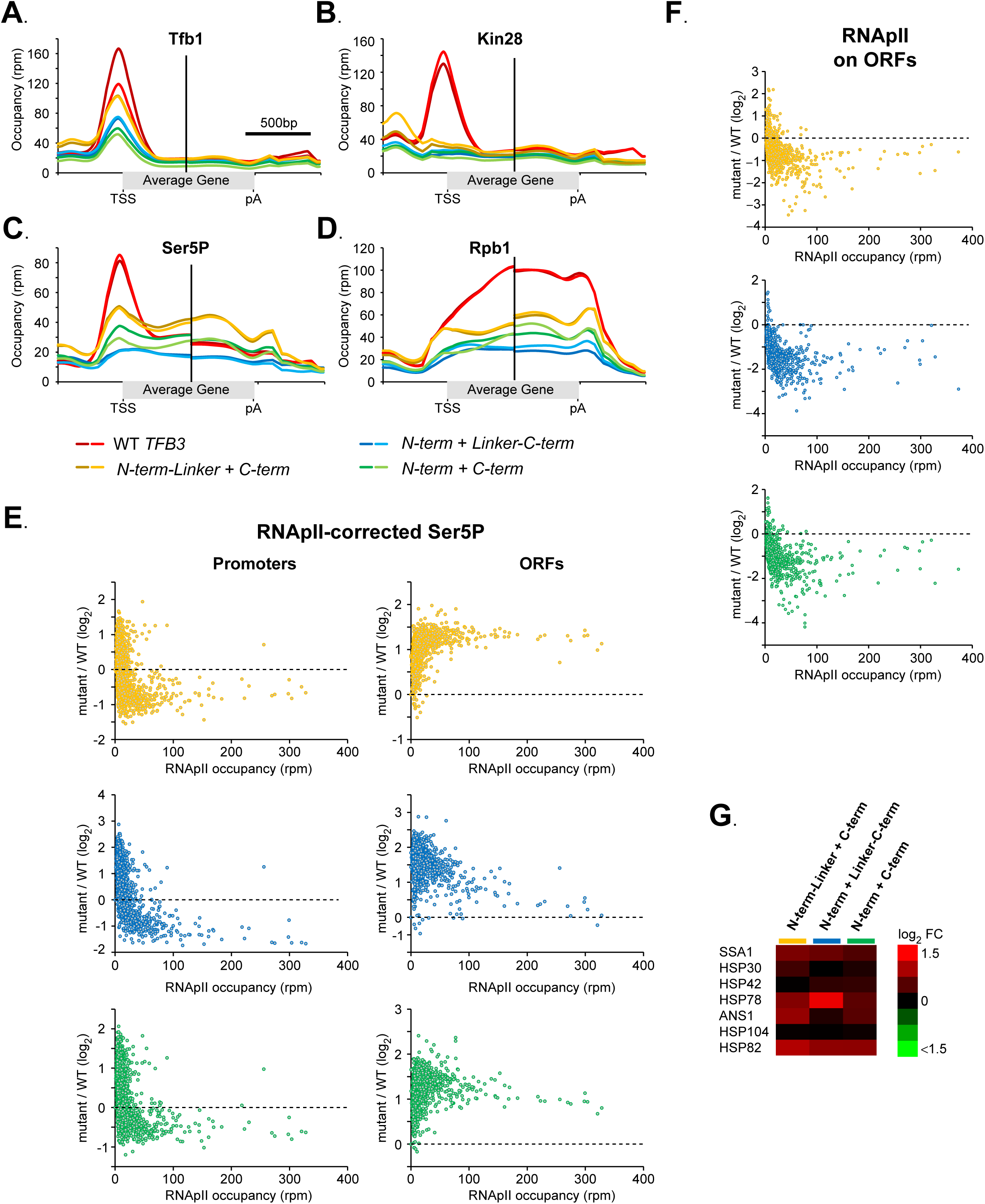
Additional ChIP-seq analysis. (**A-D**) These panels are the same metagene analysis as Figure 5A-D, but showing the individual replicates as separate lines. The Tfb3 configurations are indicated in the key at the bottom: wild-type Tfb3 in reds, N-term-Linker + C-term in yellows, N-term + Linker-C-term in blues, and N-term + C-term in greens. (**E**) For each split Tfb3 mutant (color coded as in A-D), a plot of the mutant/WT log_2_ RNApII-adjusted Ser5P ratio versus WT RNApII occupancy, for all verified genes (n=5587), over promoters (left) and ORFs (right) (see Methods). In panels E and F, the x-axis is cut at 400 rpm for clarity, excluding only three genes with higher occupancy. (**F**) Plots of the mutant/WT log_2_ ratio versus WT level for RNApII occupancy, over ORFs for all verified genes (n=5573). Yellow: N-term-Linker + C-term, Blue: N-term + Linker-C-term, and Green: N-term + C-term. (**G**) Heatmap of the mutant/WT log_2_ RNApII ratio for the seven genes that are systematically induced (mutant/WT log_2_ RNApII ratio>0) in all three split mutants. All but *ANS1* are known to be involved in stress-response.

**Figure 5 – figure supplement 2.**
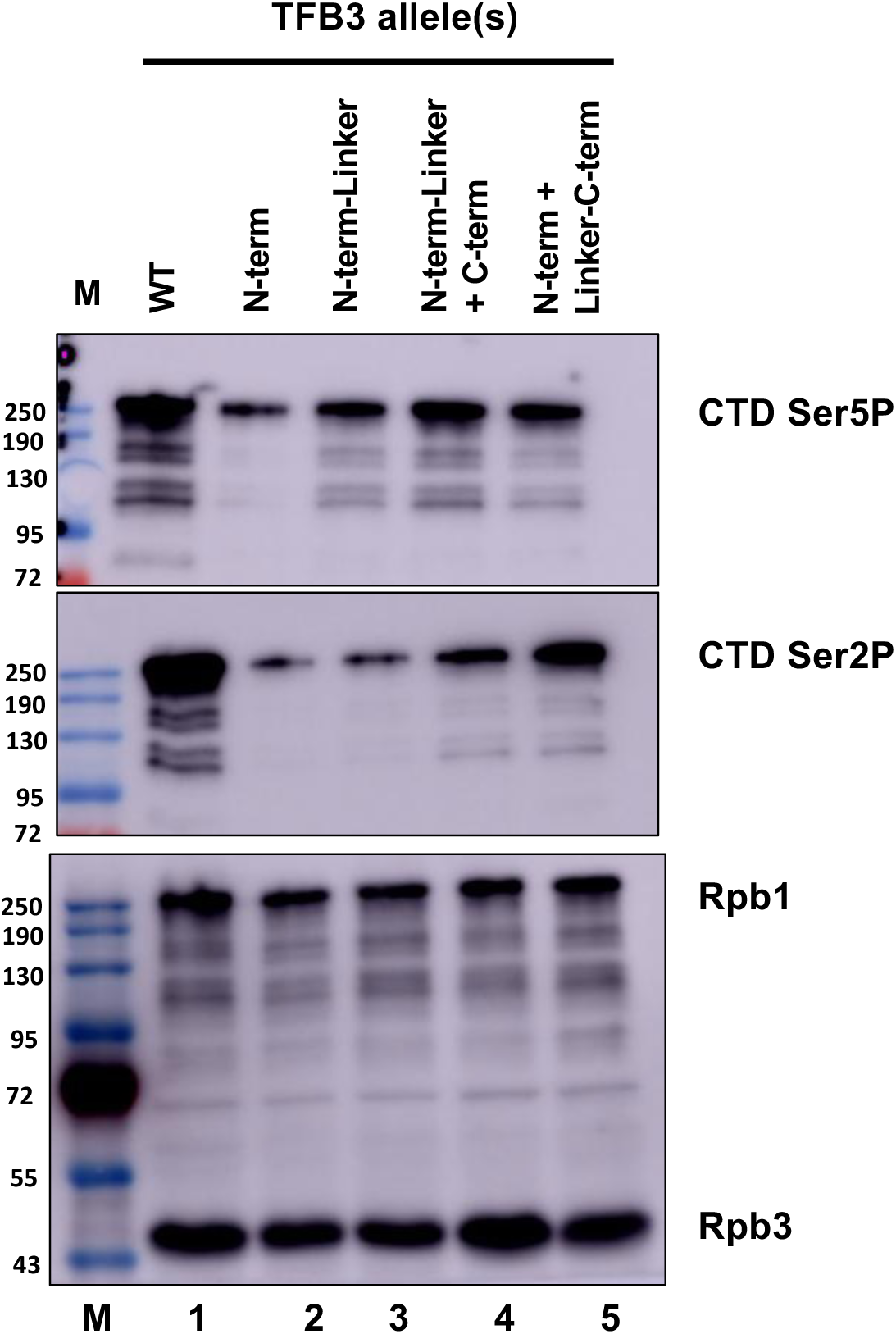
Immunoblot for CTD phosphorylation levels in strains with split Tfb3 or completely lacking the C-term. Whole cell extracts from YSB3712 (WT), YSB3710 (N-term), YSB 3711 (N-term-Linker), YSB3731 (N-term-Linker + C-term), and YSB3727 (N-term + Linker-C-term) were resolved by SDS-PAGE and immunoblotted. The top blot shows CTD Ser5P (monoclonal antibody 3E8), the middle blot shows Ser2P (monoclonal antibody 3E10), and the bottom blot shows both Rpb1 (monoclonal antibody 8WG16) and Rpb3 (monoclonal antibody 1Y26).

**Figure 5 – figure supplement 3.**
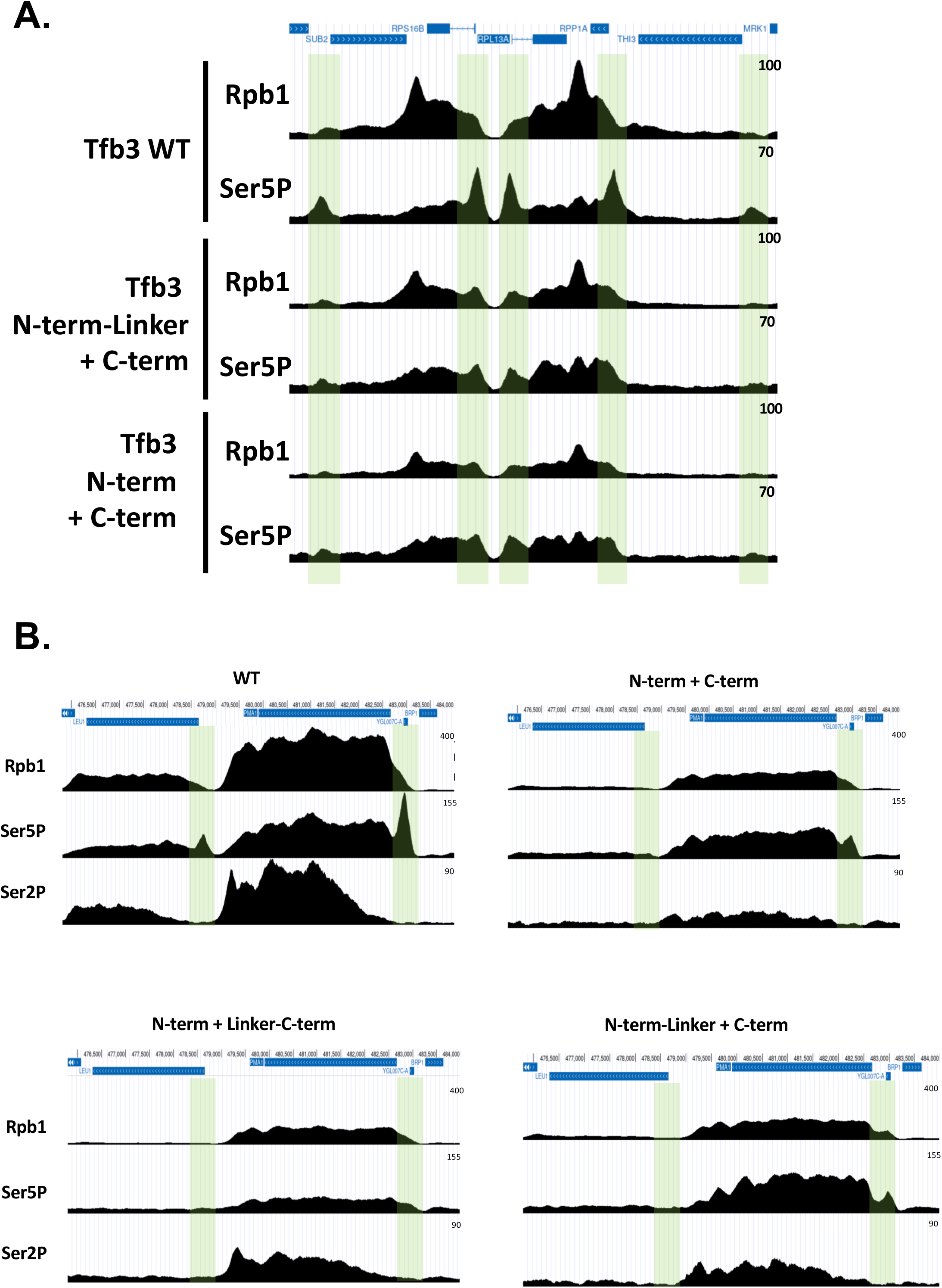
Individual gene traces from the ChIP-seq analysis show Ser5P changes in split Tfb3 strains. **A.** Rpb1 and Ser5P signals at a region containing five strong promoters (shaded in green) were plotted using the UCSC browser (https://genome.ucsc.edu). Blue bars above traces show gene locations, with arrows showing direction of transcription. Numbers in the right-hand corner show the read count scale of the y-axis. **B.** Traces of total Rpb1, Ser5P, and Ser2P at the strongly transcribed *PMA1* gene. Annotations are as in part A.

**Figure 5 - figure supplement 4.**
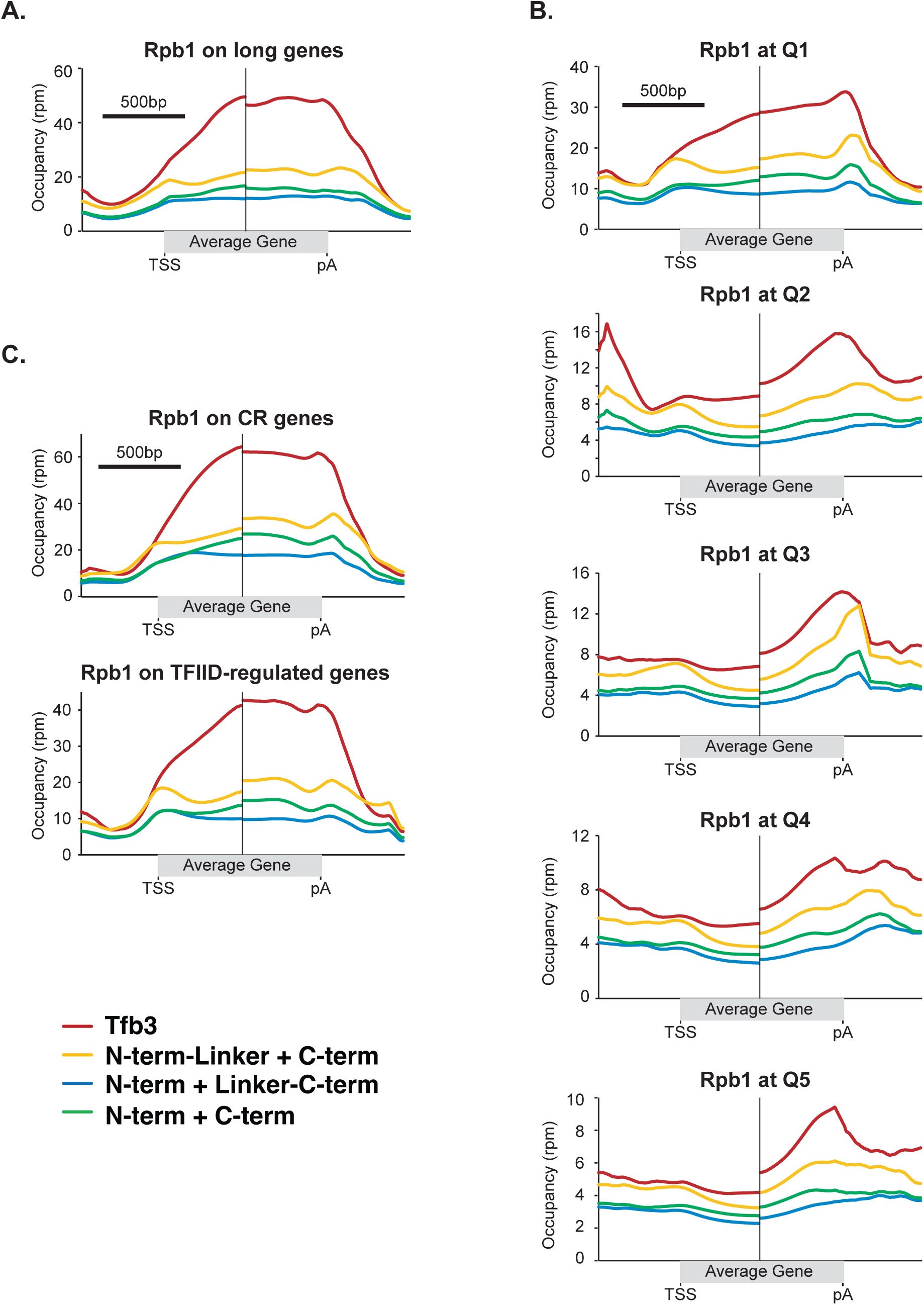
Effects of Tfb3 split mutants on RNApII occupancy are consistent across different gene classes. **A.** Metagene analysis of RNApII occupancy at “long” genes, defined as genes longer than 2 kb and having an average RNApII occupancy greater than 2 reads per million (rpm) (n=41). **B.** Metagene analysis of RNApII occupancy over all genes longer than 1 kb separated in quintiles, Q1 representing genes with the highest RNApII occupancy and Q5 the lowest. **C.** Metagene analysis of RNApII occupancy over coactivator redundant (CR) genes (top; n=151) and TFIID-regulated genes (bottom; n=55) longer then 1 kb. Gene classifications are from Donczew et al. (2020). The Tfb3 configurations are indicated in the key at the bottom: wild-type Tfb3 in red, N-term-Linker + C-term in yellow, N-term + Linker-C-term in blue, and N-term + C-term in green. To account for the differing lengths of genes, each graph shows 1 kb centered on the transcription start site (TSS) to the left of the dividing line, and 1 kb centered on the polyadenylation site (pA) to the right.

