## Supplemental Tables 1-3 for "Uncoupling the TFIIH Core and Kinase Modules Leads To Misregulated RNA Polymerase II CTD Serine 5 Phosphorylation"

**Supplemental Table S1**. Plasmids used in this study.

**Supplemental Table S2**. Yeast strains used in this study.

**Supplemental Table S3**. ChIP-Seq parameters.

**Supplemental Table 1. Plasmids used in this study.**

| Plasmid | Features | Construction or source reference |
| --- | --- | --- |
| pSBEThis7-TFB3 | T7 promoter driving N-terminal his-tagged TFB3 (1-285) [yeast TFIIH subunit], ArgU, KanR | 962 bp NcoI-Bgl II fragment bearing TFB3 open reading frame from pBS-TFB3 ORF (SB835) was cloned into NcoI + BamHI sites of pSBEThis7. Created by Michael Keogh. |
| pBS-TFB3 A | TFB3 gene with 5' flanking sequence [yeast TFIIH subunit], f1+ ori, AmpR | TFB3 gene and promoter 2.059 kb fragment PCR'd from yeast genomic DNA using primers TFB3-A (O #411) and TFB3-C (O #413) and Pfu polymerase, and blunt ligated into SrfI site of pCR-Script SK+. Created by M. Keogh. |
| pRS415-KIN28 | KIN28, LEU2, CEN/ARS, f1+ ori, AmpR | pRS426-KIN28 was digested with BamHI + HindIII. The ~1.3 kb fragment was cloned into BamHI + HindIII digested pRS415. |
| pBS-TFB3-BUR2 | TFB3 promoter driving TFB3 (1-251)-BUR2 (2-395) fusion with TFB3 5' and 3' flanking sequence, f1+ ori, AmpR | ~1.2 kb BUR2 fragment was amplified from pRS316-BUR2 (SB1209) using oligos Tfb3-Bur2 F (O #4287) and Bur2 stop Tfb3 3UTR R (O #4288). ~4.8 kb pBS-Tfb3 backbone was amplified using oligos Tfb3-3UTR F (O #4283) and Tfb3-1753 R (O #4284) on plasmid pBS-TFB3 A (SB836a). Fragments assembled through Gibson Assembly. |
| pBS-TFB3-CTK3 | TFB3 promoter driving TFB3 (1-251)-CTK3 (2-296) fusion with TFB3 5' and 3' flanking sequence, f1+ ori, AmpR | ~900 bp CTK3 fragment was amplified from pJYC4501 (F716) using oligos Tfb3-Ctk3 F (O #4285) and Ctk3 stop Tfb3 3UTR R (O #4286). ~4.8 kb pBS-Tfb3 backbone was amplified using oligos Tfb3-3UTR F (O #4283) and Tfb3-1753 R (O #4284) on plasmid pBS-TFB3 A (SB836a). Fragments assembled through Gibson Assembly. |
| pRS425-TFB3 (1-144) | TFB3 (1-144), LEU2, 2 μ ori, f1+ ori, AmpR | Oligos Tfb3- 3UTR F (O #4283) and TFB3 Ile144-Stop (O #4335) were used for inverse PCR on pRS425-TFB3 (SB1968). The resulting ~8.2 kb fragment was ligated by intramolecular Gibson isothermal assembly. |
| pRS425-TFB3 (1-251) | TFB3 (1-251), LEU2, 2 μ ori, f1+ ori, AmpR | Oligos Tfb3- 3UTR F (O #4283) and TFB3 Leu251-Stop (O #4336) were used for inverse PCR on pRS425-TFB3 (SB1968). The resulting ~8.5 kb fragment was ligated by intramolecular Gibson isothermal assembly. |
| pRS425-TFB3 | TFB3, LEU2, 2 μ ori, f1+ ori, AmpR | ~1.9 kb BamHI-Sal I fragment from pBS-TFB3 A (SB836a) cloned into the BamHI - Sal I sites of pRS425 (YV27). |
| pRS425-TFB3-MPK1 | TFB3 promoter driving TFB3 (1-251)-MPK1 (2-484) fusion with TFB3 5' and 3' flanking sequence, LEU2, 2 μ ori, f1+ ori, AmpR | ~8.5 kb pRS425-Tfb3 backbone fragment was amplified from pRS425-TFB3 (SB1968) using oligos Tfb3- 3UTR F (O #4283) and Tfb3- 1753 R (O #4284). ~1.5 kb Mpk1 fragment was amplified from genomic DNA with oligos Tfb3-Mpk1 for (O #4323) and Tfb3-Mpk1 rev (O #4324). Fragments were assembled through Gibson isothermal assembly. |
| pRS425-TFB3-BUR2 | TFB3 promoter driving TFB3 (1-251)-BUR2 (2-395) fusion with TFB3 5' and 3' flanking sequence, LEU2, 2 μ ori, f1+ ori, AmpR | ~3 kb fragment was amplified from pBS-TFB3-BUR2 (SB1973) and using oligos T3 sequencing primer (O #242) and KS poly: EcoRV-EcoRI-Pst (O #4260). ~6.8 kb pRS425 (YV27) backbone was digested with BamHI and SacI. Fragments assembled through Gibson isothermal assembly. |
| pRS425-TFB3-CTK3 | TFB3 promoter driving TFB3 (1-251)-CTK3 (2-296) fusion with TFB3 5' and 3' flanking | ~2.7 kb fragment was amplified from pBS-Tfb3-CTK3 (SB1974) using oligos T3 sequencing primer (O #242) and KS poly: EcoRV-EcoRI-Pst (O #4260). ~6.8 kb |

| | | |
|---|---|---|
| | sequence, LEU2, 2 μ ori, f1+ ori, AmpR | pRS425 (YV27) backbone was digested with BamHI and SacI. Fragments assembled through Gibson isothermal assembly. |
| pRS424-TFB3 | TFB3, TRP1, 2 μ ori, f1+ ori, AmpR | ~1.9 kb BamHI-Sal I fragment from pRS425-TFB3 (SB1968) cloned into the BamHI - Sal I sites of pRS424 (YV26). |
| pRS424-TFB3 (1-251) | TFB3 (1-251), TRP1, 2 μ ori, f1+ ori, AmpR | ~1.7 kb BamHI-SalI fragment from pRS425-TFB3 (1-251) (SB1994) and was cloned into the BamHI - Sal I sites of pRS424 (YV26). |
| pRS424-TFB3 (1-144) | TFB3 (1-144), TRP1, 2 μ ori, f1+ ori, AmpR | ~1.4 kb BamHI-SalI fragment from pRS425-TFB3 (1-144) (SB1991) was cloned into the BamHI - Sal I sites of pRS424 (YV26). |
| pRS425 | LEU2, 2 μ ori, f1+ ori, pBluescriptII SK polylinker with T7 and T3 promoters flanking, blue-white color selection, AmpR | Sikorski and Hieter (1989) Genetics 122: 19-27. |
| pRS315 | LEU2, CEN/ARS, f1+ ori, pBluescript KS+ polylinker, blue-white color selection, AmpR | Sikorski and Hieter (1989) Genetics 122: 19-27. |
| pRS425-TFB3 (1-11, 238-Stop) | TFB3 (1-11, 238-Stop), LEU2, 2 μ ori, f1+ ori, AmpR | Oligos TFB3 33 R (O #4334) and TFB3 Asp11-Pro238 (O #4339) were used for inverse PCR on pRS425-TFB3 (SB1968). The resulting ~8 kb fragment was ligated by intramolecular Gibson isothermal assembly. |
| pRS315-TFB3 (1-11, 238-Stop)-Flag1-TAP | TFB3 (1-11, 238-Stop)-Flag1-TAP, LEU2, CEN/ARS, f1+ ori, AmpR | Primers TFB3 33 R (O #4334) and TFB3 Asp11-Pro238 (O #4339) were used for inverse PCR on pSH1542 (F1255). The resulting ~7.3 kb fragment was ligated by intramolecular Gibson assembly. |
| pRS425-TFB3 (1-11, 139-Stop) | TFB3 (1-11, 139-Stop), LEU2, 2 μ ori, f1+ ori, AmpR | Primers TFB3 33 R (O #4334) and TFB3 Asp11-Leu139 (O #4337) were used for inverse PCR on pRS425-TFB3 (SB1968). The resulting ~8.3 kb fragment was ligated by intramolecular Gibson assembly. |
| pLH366 | TFB3 (Δ8-75)-Flag1-TAP, LEU2, CEN/ARS, AmpR | Warfield et al. (2016) MCB 36 (19): 2464-2475. |
| pSH1597 | TFB3, URA3, CEN/ARS, AmpR | Warfield et al. (2016) MCB 36 (19): 2464-2475. |
| pSH1542 | TFB3-Flag1-TAP, LEU2, CEN/ARS, AmpR | Warfield et al. (2016) MCB 36 (19): 2464-2475. |
| pRS315/TFB3Δ2 | TFB3Δ2 (1-275), LEU2, CEN/ARS, AmpR, f1+ ori | Feaver et al. (2000) J. Biol. Chem. 275, 5941-5946 |

**Supplemental Table 2. Yeast strains used in this study.**

| Strain | Genotype |
|---|---|
| **YSB744** | *MATα ura3-1 leu2-3,112 trp1-1 his3-11,15 kin28Δ::leu2Δ::TRP1 ade2-1 ade3-22 can1-100* [pRS426-KIN28] |
| **SHY907/ YF2456** | *MATα ura3Δ0 leu2Δ0 trp1Δ63 his3Δ200 ade2Δ::hisG lys2Δ0 met15Δ0 tfb3Δ::HygR tfb6Δ::KanMX* [pSH1597]. From Warfield et al. (2016) MCB 36 (19): 2464-2475. |
| **YSB3707** | *MATα ura3Δ0 leu2Δ0 trp1Δ63 his3Δ200 ade2Δ::hisG lys2Δ0 met15Δ0 tfb3Δ::HygR tfb6Δ::KanMX* [pSH1542] |
| **YSB3710** | *MATα ura3Δ0 leu2Δ0 trp1Δ63 his3Δ200 ade2Δ::hisG lys2Δ0 met15Δ0 tfb3Δ::HygR tfb6Δ::KanMX* [pRS425-TFB3 (1-144)] |
| **YSB3712** | *MATα ura3Δ0 leu2Δ0 trp1Δ63 his3Δ200 ade2Δ::hisG lys2Δ0 met15Δ0 tfb3Δ::HygR tfb6Δ::KanMX* [pRS425-TFB3] |
| **YSB3713** | *MATα ura3Δ0 leu2Δ0 trp1Δ63 his3Δ200 ade2Δ::hisG lys2Δ0 met15Δ0 tfb3Δ::HygR tfb6Δ::KanMX* [pRS425-TFB3-MPK1] |
| **YSB3715** | *MATα ura3Δ0 leu2Δ0 trp1Δ63 his3Δ200 ade2Δ::hisG lys2Δ0 met15Δ0 tfb3Δ::HygR tfb6Δ::KanMX* [pRS425-TFB3-BUR2] |
| **YSB3717** | *MATα ura3Δ0 leu2Δ0 trp1Δ63 his3Δ200 ade2Δ::hisG lys2Δ0 met15Δ0 tfb3Δ::HygR tfb6Δ::KanMX* [pRS425-TFB3-CTK3] |
| **YSB3722** | *MATα ura3Δ0 leu2Δ0 trp1Δ63 his3Δ200 ade2Δ::hisG lys2Δ0 met15Δ0 tfb3Δ::HygR tfb6Δ::KanMX* [pRS424-TFB3 (1-251)] |
| **YSB3704** | *MATα ura3Δ0 leu2Δ0 trp1Δ63 his3Δ200 ade2Δ::hisG lys2Δ0 met15Δ0 tfb3Δ::HygR tfb6Δ::KanMX* [pRS424-TFB3 (1-144)] |
| **YSB3723** | *MATα ura3Δ0 leu2Δ0 trp1Δ63 his3Δ200 ade2Δ::hisG lys2Δ0 met15Δ0 tfb3Δ::HygR tfb6Δ::KanMX* [pRS424-TFB3 (1-144), pSH1542] |
| **YSB3724** | *MATα ura3Δ0 leu2Δ0 trp1Δ63 his3Δ200 ade2Δ::hisG lys2Δ0 met15Δ0 tfb3Δ::HygR tfb6Δ::KanMX* [pRS424-TFB3 (1-144), pRS425] |
| **YSB3725** | *MATα ura3Δ0 leu2Δ0 trp1Δ63 his3Δ200 ade2Δ::hisG lys2Δ0 met15Δ0 tfb3Δ::HygR tfb6Δ::KanMX* [pRS424-TFB3 (1-144), pRS425-TFB3 (1-11, 238-Stop)] |
| **YSB3726** | *MATα ura3Δ0 leu2Δ0 trp1Δ63 his3Δ200 ade2Δ::hisG lys2Δ0 met15Δ0 tfb3Δ::HygR tfb6Δ::KanMX* [pRS424-TFB3 (1-144), pRS315-TFB3 (1-11, 238-Stop)-Flag1-TAP] |
| **YSB3727** | *MATα ura3Δ0 leu2Δ0 trp1Δ63 his3Δ200 ade2Δ::hisG lys2Δ0 met15Δ0 tfb3Δ::HygR tfb6Δ::KanMX* [pRS424-TFB3 (1-144), pRS425-TFB3 (1-11, 139-Stop)] |
| **YSB3728** | *MATα ura3Δ0 leu2Δ0 trp1Δ63 his3Δ200 ade2Δ::hisG lys2Δ0 met15Δ0 tfb3Δ::HygR tfb6Δ::KanMX* [pRS424-TFB3 (1-144), pLH366] |
| **YSB3729** | *MATα ura3Δ0 leu2Δ0 trp1Δ63 his3Δ200 ade2Δ::hisG lys2Δ0 met15Δ0 tfb3Δ::HygR tfb6Δ::KanMX* [pRS424-TFB3 (1-251), pSH1542] |
| **YSB3730** | *MATα ura3Δ0 leu2Δ0 trp1Δ63 his3Δ200 ade2Δ::hisG lys2Δ0 met15Δ0 tfb3Δ::HygR tfb6Δ::KanMX* [pRS424-TFB3 (1-251), pRS425] |
| **YSB3731** | *MATα ura3Δ0 leu2Δ0 trp1Δ63 his3Δ200 ade2Δ::hisG lys2Δ0 met15Δ0 tfb3Δ::HygR tfb6Δ::KanMX* [pRS424-TFB3 (1-251), pRS425-TFB3 (1-11, 238-Stop)] |
| **YSB3732** | *MATα ura3Δ0 leu2Δ0 trp1Δ63 his3Δ200 ade2Δ::hisG lys2Δ0 met15Δ0 tfb3Δ::HygR tfb6Δ::KanMX* [pRS424-TFB3 (1-251), pRS315-TFB3 (1-11, 238-Stop)-Flag1-TAP] |
| **YSB3733** | *MATα ura3Δ0 leu2Δ0 trp1Δ63 his3Δ200 ade2Δ::hisG lys2Δ0 met15Δ0 tfb3Δ::HygR tfb6Δ::KanMX* [pRS424-TFB3 (1-251), pRS425-TFB3 (1-11, 139-Stop)] |
| **YSB3734** | *MATα ura3Δ0 leu2Δ0 trp1Δ63 his3Δ200 ade2Δ::hisG lys2Δ0 met15Δ0 tfb3Δ::HygR tfb6Δ::KanMX* [pRS424-TFB3 (1-251), pLH366] |
| **YSB3786** | *MATα ura3-1 leu2-3,112 trp1-1 his3-11,15 kin28Δ::leu2Δ::TRP1 ade2-1 ade3-22 can1-100* [pRS415-KIN28] |
| **YSB3787** | *MATα ura3Δ0 leu2Δ0 trp1Δ63 his3Δ200 ade2Δ::hisG lys2Δ0 met15Δ0 tfb3Δ::HygR tfb6Δ::KanMX* [pRS424-TFB3] |
| **YSB3788** | *MATα ura3Δ0 leu2Δ0 trp1Δ63 his3Δ200 ade2Δ::hisG lys2Δ0 met15Δ0 tfb3Δ::HygR tfb6Δ::KanMX* [pRS424-TFB3, pRS425] |
| **YSB207** | *MATa ura3-52 leu2-3,112 his3Δ200 tfb1Δ::LEU2* [pRS316-TFB1] From Matsui et al., (1995) |

| | |
|---|---|
| | Nucleic Acids Res 23, 767-772. |
| **YSB260** | *MATa ura3-52 leu2-3,112 his3Δ200 tfb1Δ::LEU2* [pRS313-tfb1-101] From Matsui et al., (1995) Nucleic Acids Res 23, 767-772. |

### Supplemental Table S3a: ChIP-seq parameters for S. cerevisiae reads

| Sample | Total reads | Mapped reads | % Mapped reads | Deduplicated reads | % Deduplicated reads | Fragments average size | Fragments size std | Pearson correlation |
|---|---|---|---|---|---|---|---|---|
| CJ1_ChIPseq_Input_Tfb3WT_Spike_R1_20221206 | 28060948 | 24316268 | 86.66% | 17666200 | 62.96% | 187.151693 | 54.84188419 | |
| CJ2_ChIPseq_Input_Tfb3WT_Spike_R2_20221206 | 25142766 | 21683658 | 86.24% | 16285104 | 64.77% | 183.7562841 | 52.72211465 | |
| CJ3_ChIPseq_Input_tfb3N-LC_Spike_R1_20221206 | 30338000 | 28318312 | 93.34% | 19901330 | 65.60% | 182.0606635 | 51.85993552 | |
| CJ4_ChIPseq_Input_tfb3N-LC_Spike_R2_20221206 | 25623514 | 23744682 | 92.67% | 18112754 | 70.69% | 193.12372 | 59.31659928 | |
| CJ5_ChIPseq_Input_tfb3N-C_Spike_R1_20221206 | 26168988 | 23924892 | 91.42% | 18380502 | 70.24% | 185.0257088 | 53.5344394 | |
| CJ6_ChIPseq_Input_tfb3N-C_Spike_R2_20221206 | 28575464 | 26292900 | 92.01% | 20354770 | 71.23% | 192.773337 | 58.00570866 | |
| CJ7_ChIPseq_Input_tfb3NL-C_Spike_R1_20221206 | 31451666 | 27197436 | 86.47% | 19478246 | 61.93% | 185.1408434 | 54.21921369 | |
| CJ8_ChIPseq_Input_tfb3NL-C_Spike_R2_20221206 | 27660176 | 24037478 | 86.90% | 17810316 | 64.39% | 187.0205919 | 55.0058291 | |
| CJ9_ChIPseq_IP_Tfb1_Tfb3WT_Spike_R1_20221206 | 21933424 | 18706082 | 85.29% | 8523750 | 38.86% | 178.9884375 | 50.4586975 | |
| CJ11_ChIPseq_IP_Tfb1_Tfb3WT_Spike_R2_20221206 | 31388434 | 27114844 | 86.38% | 15064408 | 47.99% | 180.6129835 | 50.89617922 | |
| CJ10_ChIPseq_IP_Kin28_Tfb3WT_Spike_R1_20221206 | 25212688 | 22084698 | 87.59% | 11771758 | 46.69% | 183.0545385 | 53.52558161 | |
| CJ12_ChIPseq_IP_Kin28_Tfb3WT_Spike_R2_20221206 | 27504962 | 24137616 | 87.76% | 12353214 | 44.91% | 187.8508748 | 55.94383818 | |
| CJ13_ChIPseq_IP_Tfb1_tfb3N-LC_Spike_R1_20221206 | 26079084 | 24104222 | 92.43% | 14991702 | 57.49% | 186.3978388 | 54.85202601 | |
| CJ15_ChIPseq_IP_Tfb1_tfb3N-LC_Spike_R2_20221206 | 28492222 | 26201452 | 91.96% | 16217586 | 56.92% | 184.8119941 | 53.90919534 | |
| CJ14_ChIPseq_IP_Kin28_tfb3N-LC_Spike_R1_20221206 | 26720222 | 24929216 | 93.30% | 15482036 | 57.94% | 186.9584852 | 55.33301124 | |
| CJ16_ChIPseq_IP_Kin28_tfb3N-LC_Spike_R2_20221206 | 30556826 | 28384914 | 92.89% | 16973328 | 55.55% | 191.2947476 | 56.93516834 | |
| CJ17_ChIPseq_IP_Tfb1_tfb3N-C_Spike_R1_20221206 | 28014174 | 25070362 | 89.49% | 12797216 | 45.68% | 186.2883061 | 55.37930129 | |
| CJ19_ChIPseq_IP_Tfb1_tfb3N-C_Spike_R2_20221206 | 23685496 | 20727300 | 87.51% | 7118384 | 30.05% | 181.9331486 | 52.28734443 | |
| CJ18_ChIPseq_IP_Kin28_tfb3N-C_Spike_R1_20221206 | 26253552 | 23887458 | 90.99% | 13124260 | 49.99% | 190.0250035 | 55.02351744 | |
| CJ20_ChIPseq_IP_Kin28_tfb3N-C_Spike_R2_20221206 | 27987324 | 25315522 | 90.45% | 10788264 | 38.55% | 186.4347784 | 54.96583003 | |
| CJ21_ChIPseq_IP_Tfb1_tfb3NL-C_Spike_R1_20221206 | 21097654 | 17251128 | 81.77% | 5973408 | 28.31% | 184.6293489 | 52.40626339 | |
| CJ23_ChIPseq_IP_Tfb1_tfb3NL-C_Spike_R2_20221206 | 25005018 | 21356392 | 85.41% | 7346354 | 29.38% | 182.4302594 | 52.97200481 | |
| CJ22_ChIPseq_IP_Kin28_tfb3NL-C_Spike_R1_20221206 | 22756574 | 19667140 | 86.42% | 10115178 | 44.45% | 181.6662182 | 52.00454101 | |
| CJ24_ChIPseq_IP_Kin28_tfb3NL-C_Spike_R2_20221206 | 24373472 | 21125784 | 86.68% | 8198594 | 33.64% | 197.0143442 | 60.81813163 | |
| CJ25_ChIPseq_IP_8WG16_Tfb3WT_Spike_R1_20221206 | 27441896 | 23822720 | 86.81% | 18410874 | 67.09% | 193.5061943 | 56.02529553 | |
| CJ28_ChIPseq_IP_8WG16_Tfb3WT_Spike_R2_20221206 | 25996340 | 22584706 | 86.88% | 17075896 | 65.69% | 194.006966 | 55.81883403 | |
| CJ26_ChIPseq_IP_3E8_Tfb3WT_Spike_R1_20221206 | 25838296 | 21654764 | 83.81% | 14876982 | 57.58% | 184.3699006 | 52.06012194 | |
| CJ29_ChIPseq_IP_3E8_Tfb3WT_Spike_R2_20221206 | 28040716 | 21924548 | 78.19% | 15023372 | 53.58% | 183.4677026 | 52.19602674 | |
| CJ31_ChIPseq_IP_8WG16_tfb3N-LC_Spike_R1_20221206 | 30495908 | 27080862 | 88.80% | 18984972 | 62.25% | 189.7476167 | 55.15932956 | |
| CJ34_ChIPseq_IP_8WG16_tfb3N-LC_Spike_R2_20221206 | 25196604 | 22133444 | 87.84% | 16020930 | 63.58% | 189.3908446 | 54.63351052 | |
| CJ32_ChIPseq_IP_3E8_tfb3N-LC_Spike_R1_20221206 | 27969570 | 25873014 | 92.50% | 16869978 | 60.32% | 191.6410608 | 55.85065663 | |
| CJ35_ChIPseq_IP_3E8_tfb3N-LC_Spike_R2_20221206 | 22792800 | 20375434 | 89.39% | 13537306 | 59.39% | 186.0942068 | 53.96482432 | |
| CJ37_ChIPseq_IP_8WG16_tfb3N-C_Spike_R1_20221206 | 27586338 | 24259098 | 87.94% | 17339088 | 62.85% | 185.0421983 | 51.76648932 | |
| CJ40_ChIPseq_IP_8WG16_tfb3N-C_Spike_R2_20221206 | 23630278 | 20684686 | 87.53% | 14216836 | 60.16% | 192.4918004 | 53.96706389 | |
| CJ38_ChIPseq_IP_3E8_tfb3N-C_Spike_R1_20221206 | 25817692 | 22883660 | 88.64% | 14284602 | 55.33% | 181.8681642 | 51.2903419 | |
| CJ41_ChIPseq_IP_3E8_tfb3N-C_Spike_R2_20221206 | 22984208 | 20364348 | 88.60% | 10643430 | 46.31% | 179.9475536 | 48.44600528 | |
| CJ43_ChIPseq_IP_8WG16_tfb3NL-C_Spike_R1_20221206 | 31686578 | 26214330 | 82.73% | 18765146 | 59.22% | 198.0127415 | 55.85851478 | |
| CJ46_ChIPseq_IP_8WG16_tfb3NL-C_Spike_R2_20221206 | 28413026 | 23563850 | 82.93% | 16894050 | 59.46% | 199.4182276 | 56.07545824 | |
| CJ44_ChIPseq_IP_3E8_tfb3NL-C_Spike_R1_20221206 | 22729102 | 18672918 | 82.15% | 10778412 | 47.42% | 194.0807191 | 56.0081025 | |
| CJ47_ChIPseq_IP_3E8_tfb3NL-C_Spike_R2_20221206 | 27089408 | 22290936 | 82.29% | 12664346 | 46.75% | 190.8298633 | 54.2312006 | |
| ChIPseq_Input_Tfb3WT_Spike_CJ1-CJ2_20221206 | | | | 33951304 | | 185.523049 | 53.86224722 | 0.9987 |
| ChIPseq_Input_tfb3N-LC_Spike_CJ3-CJ4_20221206 | | | | 38014084 | | 187.3319314 | 55.81203566 | 0.9929 |
| ChIPseq_Input_tfb3N-C_Spike_CJ5-CJ6_20221206 | | | | 38735272 | | 189.0969643 | 56.06225203 | 0.9982 |
| ChIPseq_Input_tfb3NL-C_Spike_CJ7-CJ8_20221206 | | | | 37288562 | | 186.0386767 | 54.60441417 | 0.9918 |
| ChIPseq_IP_Tfb1_Tfb3WT_Spike_CJ9-CJ11_20221206 | | | | 23588158 | | 180.0259422 | 50.74452676 | 0.9937 |
| ChIPseq_IP_Kin28_Tfb3WT_Spike_CJ10-CJ12_20221206 | | | | 24124972 | | 185.5105069 | 54.82962941 | 0.9994 |
| ChIPseq_IP_Tfb1_tfb3N-LC_Spike_CJ13-CJ15_20221206 | | | | 31209288 | | 185.5737709 | 54.36990612 | 0.9955 |
| ChIPseq_IP_Kin28_tfb3N-LC_Spike_CJ14-CJ16_20221206 | | | | 32455364 | | 189.2262399 | 56.21833234 | 0.9948 |
| ChIPseq_IP_Tfb1_tfb3N-C_Spike_CJ17-CJ19_20221206 | | | | 19915600 | | 184.7316529 | 54.334475 | 0.988 |
| ChIPseq_IP_Kin28_tfb3N-C_Spike_CJ18-CJ20_20221206 | | | | 23912524 | | 188.4052541 | 55.02650556 | 0.9945 |
| ChIPseq_IP_Tfb1_tfb3NL-C_Spike_CJ21-CJ23_20221206 | | | | 13319762 | | 183.4164677 | 52.73038151 | 0.9976 |
| ChIPseq_IP_Kin28_tfb3NL-C_Spike_CJ22-CJ24_20221206 | | | | 18313772 | | 188.5371705 | 56.63808578 | 0.9967 |
| ChIPseq_IP_8WG16_Tfb3WT_Spike_CJ25-CJ28_20221206 | | | | 35486770 | | 193.7471609 | 55.92660148 | 0.9982 |
| ChIPseq_IP_3E8_Tfb3WT_Spike_CJ26-CJ29_20221206 | | | | 29900354 | | 183.916593 | 52.13040131 | 0.9802 |
| ChIPseq_IP_8WG16_tfb3N-LC_Spike_CJ31-CJ34_20221206 | | | | 35005902 | | 189.5843351 | 54.91959214 | 0.9305 |
| ChIPseq_IP_3E8_tfb3N-LC_Spike_CJ32-CJ35_20221206 | | | | 30407284 | | 189.1716045 | 55.08808583 | 0.9763 |
| ChIPseq_IP_8WG16_tfb3N-C_Spike_CJ37-CJ40_20221206 | | | | 31555924 | | 188.3984547 | 52.89928193 | 0.9869 |
| ChIPseq_IP_3E8_tfb3N-C_Spike_CJ38-CJ41_20221206 | | | | 24928032 | | 181.0481281 | 50.10467126 | 0.9863 |
| ChIPseq_IP_8WG16_tfb3NL-C_Spike_CJ43-CJ46_20221206 | | | | 35659196 | | 198.6786105 | 55.96579816 | 0.985 |
| ChIPseq_IP_3E8_tfb3NL-C_Spike_CJ44-CJ47_20221206 | | | | 23442758 | | 192.324528 | 55.07913115 | 0.9835 |

**Supplemental Table S3b: ChIP-seq parameters for S. pombe reads**

| Sample | Total reads | Mapped reads | % Mapped reads | Deduplicated reads | % Deduplicated reads | Fragments average size | Fragments size std |
|---|---|---|---|---|---|---|---|
| CJ1_ChIPseq_Input_Tfb3WT_Spike_R1_20221206 | 28060948 | 846652 | 3.02% | 623366 | 2.22% | 185.1013305 | 53.83830897 |
| CJ2_ChIPseq_Input_Tfb3WT_Spike_R2_20221206 | 25142766 | 781614 | 3.11% | 596604 | 2.37% | 181.5391013 | 51.55880425 |
| CJ3_ChIPseq_Input_tfb3N-LC_Spike_R1_20221206 | 30338000 | 695654 | 2.29% | 492432 | 1.62% | 179.9999756 | 50.50852705 |
| CJ4_ChIPseq_Input_tfb3N-LC_Spike_R2_20221206 | 25623514 | 560076 | 2.19% | 428710 | 1.67% | 191.1615171 | 58.56076855 |
| CJ5_ChIPseq_Input_tfb3N-C_Spike_R1_20221206 | 26168988 | 670838 | 2.56% | 519310 | 1.98% | 182.5892319 | 52.50833339 |
| CJ6_ChIPseq_Input_tfb3N-C_Spike_R2_20221206 | 28575464 | 645568 | 2.26% | 503894 | 1.76% | 189.4855466 | 56.74773664 |
| CJ7_ChIPseq_Input_tfb3NL-C_Spike_R1_20221206 | 31451666 | 1368696 | 4.35% | 990526 | 3.15% | 183.7836241 | 53.83598574 |
| CJ8_ChIPseq_Input_tfb3NL-C_Spike_R2_20221206 | 27660176 | 1332618 | 4.82% | 995828 | 3.60% | 185.958535 | 54.71051443 |
| CJ9_ChIPseq_IP_Tfb1_Tfb3WT_Spike_R1_20221206 | 21933424 | 732152 | 3.34% | 335374 | 1.53% | 176.9659902 | 49.48895332 |
| CJ11_ChIPseq_IP_Tfb1_Tfb3WT_Spike_R2_20221206 | 31388434 | 1070344 | 3.41% | 619132 | 1.97% | 177.9171195 | 49.28553322 |
| CJ10_ChIPseq_IP_Kin28_Tfb3WT_Spike_R1_20221206 | 25212688 | 674476 | 2.68% | 379710 | 1.51% | 179.4113244 | 51.7300538 |
| CJ12_ChIPseq_IP_Kin28_Tfb3WT_Spike_R2_20221206 | 27504962 | 747658 | 2.72% | 408752 | 1.49% | 184.1809263 | 54.14960845 |
| CJ13_ChIPseq_IP_Tfb1_tfb3N-LC_Spike_R1_20221206 | 26079084 | 728480 | 2.79% | 457532 | 1.75% | 183.5296854 | 53.51132098 |
| CJ15_ChIPseq_IP_Tfb1_tfb3N-LC_Spike_R2_20221206 | 28492222 | 790138 | 2.77% | 494990 | 1.74% | 181.7431908 | 52.49740535 |
| CJ14_ChIPseq_IP_Kin28_tfb3N-LC_Spike_R1_20221206 | 26720222 | 666442 | 2.49% | 422284 | 1.58% | 183.0614989 | 53.28236879 |
| CJ16_ChIPseq_IP_Kin28_tfb3N-LC_Spike_R2_20221206 | 30556826 | 719998 | 2.36% | 434818 | 1.42% | 187.2660699 | 55.15772767 |
| CJ17_ChIPseq_IP_Tfb1_tfb3N-C_Spike_R1_20221206 | 28014174 | 1063794 | 3.80% | 544726 | 1.94% | 182.2505076 | 53.30852239 |
| CJ19_ChIPseq_IP_Tfb1_tfb3N-C_Spike_R2_20221206 | 23685496 | 1072190 | 4.53% | 362356 | 1.53% | 178.0362296 | 50.24777303 |
| CJ18_ChIPseq_IP_Kin28_tfb3N-C_Spike_R1_20221206 | 26253552 | 767778 | 2.92% | 433724 | 1.65% | 185.1550525 | 53.05492754 |
| CJ20_ChIPseq_IP_Kin28_tfb3N-C_Spike_R2_20221206 | 27987324 | 834902 | 2.98% | 372238 | 1.33% | 180.7236016 | 51.9326008 |
| CJ21_ChIPseq_IP_Tfb1_tfb3NL-C_Spike_R1_20221206 | 21097654 | 1141534 | 5.41% | 378344 | 1.79% | 182.2998065 | 51.77239721 |
| CJ23_ChIPseq_IP_Tfb1_tfb3NL-C_Spike_R2_20221206 | 25005018 | 1487086 | 5.95% | 495676 | 1.98% | 180.1907738 | 52.38100476 |
| CJ22_ChIPseq_IP_Kin28_tfb3NL-C_Spike_R1_20221206 | 22756574 | 1136130 | 4.99% | 610830 | 2.68% | 177.3449569 | 49.83973602 |
| CJ24_ChIPseq_IP_Kin28_tfb3NL-C_Spike_R2_20221206 | 24373472 | 1178548 | 4.84% | 458122 | 1.88% | 192.6899821 | 58.951525 |
| CJ25_ChIPseq_IP_8WG16_Tfb3WT_Spike_R1_20221206 | 27441896 | 1128564 | 4.11% | 884000 | 3.22% | 190.3345271 | 54.68234901 |
| CJ28_ChIPseq_IP_8WG16_Tfb3WT_Spike_R2_20221206 | 25996340 | 1092832 | 4.20% | 835682 | 3.21% | 190.8983728 | 54.47781093 |
| CJ26_ChIPseq_IP_3E8_Tfb3WT_Spike_R1_20221206 | 25838296 | 1662118 | 6.43% | 1154128 | 4.47% | 181.3311574 | 50.62594455 |
| CJ29_ChIPseq_IP_3E8_Tfb3WT_Spike_R2_20221206 | 28040716 | 1958358 | 6.98% | 1357596 | 4.84% | 180.5729215 | 50.74377243 |
| CJ27_ChIPseq_IP_3E10_Tfb3WT_Spike_R1_20221206 | 30278142 | 3027838 | 10.00% | 507444 | 1.68% | 180.0213304 | 51.4626712 |
| CJ30_ChIPseq_IP_3E10_Tfb3WT_Spike_R2_20221206 | 31671338 | 3343260 | 10.56% | 484610 | 1.53% | 178.9929964 | 52.28392339 |
| CJ31_ChIPseq_IP_8WG16_tfb3N-LC_Spike_R1_20221206 | 30495908 | 2113824 | 6.93% | 1484396 | 4.87% | 185.3269855 | 53.05985514 |
| CJ34_ChIPseq_IP_8WG16_tfb3N-LC_Spike_R2_20221206 | 25196604 | 1773960 | 7.04% | 1293002 | 5.13% | 185.4773775 | 52.69852257 |
| CJ32_ChIPseq_IP_3E8_tfb3N-LC_Spike_R1_20221206 | 27969570 | 1003486 | 3.59% | 657046 | 2.35% | 187.0917653 | 53.92475137 |
| CJ35_ChIPseq_IP_3E8_tfb3N-LC_Spike_R2_20221206 | 22792800 | 1243894 | 5.46% | 824336 | 3.62% | 181.9736928 | 51.7804567 |
| CJ33_ChIPseq_IP_3E10_tfb3N-LC_Spike_R1_20221206 | 42940988 | 1247534 | 2.91% | 627904 | 1.46% | 183.4230264 | 52.97077163 |
| CJ36_ChIPseq_IP_3E10_tfb3N-LC_Spike_R2_20221206 | 36916590 | 1791252 | 4.85% | 508930 | 1.38% | 186.0975183 | 56.74859061 |
| CJ37_ChIPseq_IP_8WG16_tfb3N-C_Spike_R1_20221206 | 27586338 | 1659278 | 6.01% | 1199388 | 4.35% | 180.2060768 | 49.49942587 |
| CJ40_ChIPseq_IP_8WG16_tfb3N-C_Spike_R2_20221206 | 23630278 | 1365400 | 5.78% | 950242 | 4.02% | 187.1708954 | 51.92034006 |
| CJ38_ChIPseq_IP_3E8_tfb3N-C_Spike_R1_20221206 | 25817692 | 1314274 | 5.09% | 820672 | 3.18% | 177.7428912 | 49.17595847 |
| CJ41_ChIPseq_IP_3E8_tfb3N-C_Spike_R2_20221206 | 22984208 | 1090638 | 4.75% | 568434 | 2.47% | 175.8922302 | 46.61382335 |
| CJ39_ChIPseq_IP_3E10_tfb3N-C_Spike_R1_20221206 | 51180370 | 2894724 | 5.66% | 254696 | 0.50% | 179.5519757 | 51.41823937 |
| CJ42_ChIPseq_IP_3E10_tfb3N-C_Spike_R2_20221206 | 34504488 | 1583728 | 4.59% | 258710 | 0.75% | 183.0624792 | 52.30351672 |
| CJ43_ChIPseq_IP_8WG16_tfb3NL-C_Spike_R1_20221206 | 31686578 | 2766612 | 8.73% | 2016986 | 6.37% | 194.0235222 | 54.1382237 |
| CJ46_ChIPseq_IP_8WG16_tfb3NL-C_Spike_R2_20221206 | 28413026 | 2677376 | 9.42% | 1948088 | 6.86% | 195.6566202 | 54.61740573 |
| CJ44_ChIPseq_IP_3E8_tfb3NL-C_Spike_R1_20221206 | 22729102 | 1896804 | 8.35% | 1092670 | 4.81% | 190.4241445 | 54.41026838 |
| CJ47_ChIPseq_IP_3E8_tfb3NL-C_Spike_R2_20221206 | 27089408 | 2631848 | 9.72% | 1495020 | 5.52% | 187.6967934 | 52.80320334 |
| CJ45_ChIPseq_IP_3E10_tfb3NL-C_Spike_R1_20221206 | 30375814 | 2941948 | 9.69% | 455950 | 1.50% | 181.3833798 | 54.21909823 |
| CJ48_ChIPseq_IP_3E10_tfb3NL-C_Spike_R2_20221206 | 31921716 | 4194896 | 13.14% | 340390 | 1.07% | 181.5553042 | 51.34714194 |
| ChIPseq_Input_Tfb3WT_Spike_CJ1-CJ2_20221206 | | | | 1219970 | | 183.3592875 | 52.76588264 |
| ChIPseq_Input_tfb3N-LC_Spike_CJ3-CJ4_20221206 | | | | 921142 | | 185.1946844 | 54.68865897 |
| ChIPseq_Input_tfb3N-C_Spike_CJ5-CJ6_20221206 | | | | 1023204 | | 185.9854379 | 54.74584347 |
| ChIPseq_Input_tfb3NL-C_Spike_CJ7-CJ8_20221206 | | | | 1986354 | | 184.8739822 | 54.28704405 |
| ChIPseq_IP_Tfb1_Tfb3WT_Spike_CJ9-CJ11_20221206 | | | | 954506 | | 177.5829319 | 49.3591389 |
| ChIPseq_IP_Kin28_Tfb3WT_Spike_CJ10-CJ12_20221206 | | | | 788462 | | 181.8839665 | 53.0516712 |
| ChIPseq_IP_Tfb1_tfb3N-LC_Spike_CJ13-CJ15_20221206 | | | | 952522 | | 182.601311 | 52.99430978 |
| ChIPseq_IP_Kin28_tfb3N-LC_Spike_CJ14-CJ16_20221206 | | | | 857102 | | 185.1945276 | 54.282517 |
| ChIPseq_IP_Tfb1_tfb3N-C_Spike_CJ17-CJ19_20221206 | | | | 907082 | | 180.5670116 | 52.14821133 |
| ChIPseq_IP_Kin28_tfb3N-C_Spike_CJ18-CJ20_20221206 | | | | 805962 | | 183.1083624 | 52.58591834 |
| ChIPseq_IP_Tfb1_tfb3NL-C_Spike_CJ21-CJ23_20221206 | | | | 874020 | | 181.1037276 | 52.12883967 |
| ChIPseq_IP_Kin28_tfb3NL-C_Spike_CJ22-CJ24_20221206 | | | | 1068952 | | 183.9213922 | 54.46554015 |
| ChIPseq_IP_8WG16_Tfb3WT_Spike_CJ25-CJ28_20221206 | | | | 1719682 | | 190.6085288 | 54.58374492 |
| ChIPseq_IP_3E8_Tfb3WT_Spike_CJ26-CJ29_20221206 | | | | 2511724 | | 180.9213281 | 50.69105321 |
| ChIPseq_IP_3E10_Tfb3WT_Spike_CJ27-CJ30_20221206 | | | | 992054 | | 179.518998 | 51.86796548 |
| ChIPseq_IP_8WG16_tfb3N-LC_Spike_CJ31-CJ34_20221206 | | | | 2777398 | | 185.3969996 | 52.89198006 |
| ChIPseq_IP_3E8_tfb3N-LC_Spike_CJ32-CJ35_20221206 | | | | 1481382 | | 184.2437413 | 52.80350633 |
| ChIPseq_IP_3E10_tfb3N-LC_Spike_CJ33-CJ36_20221206 | | | | 1136834 | | 184.6203245 | 54.7103866 |
| ChIPseq_IP_8WG16_tfb3N-C_Spike_CJ37-CJ40_20221206 | | | | 2149630 | | 183.2848686 | 50.70197932 |
| ChIPseq_IP_3E8_tfb3N-C_Spike_CJ38-CJ41_20221206 | | | | 1389106 | | 176.985585 | 48.15256033 |
| ChIPseq_IP_3E10_tfb3N-C_Spike_CJ39-CJ42_20221206 | | | | 513406 | | 181.3209507 | 51.89581686 |
| ChIPseq_IP_8WG16_tfb3NL-C_Spike_CJ43-CJ46_20221206 | | | | 3965074 | | 194.8258827 | 54.38029447 |
| ChIPseq_IP_3E8_tfb3NL-C_Spike_CJ44-CJ47_20221206 | | | | 2587690 | | 188.8484363 | 53.50462746 |
| ChIPseq_IP_3E10_tfb3NL-C_Spike_CJ45-CJ48_20221206 | | | | 796340 | | 181.4568677 | 53.01054631 |
